# Chlorosis as a developmental program in cyanobacteria: the proteomic fundament for survival and awakening

**DOI:** 10.1101/325761

**Authors:** Philipp Spät, Alexander Klotz, Sascha Rexroth, Boris Maček, Karl Forchhammer

## Abstract

Cyanobacteria that do not fix atmospheric nitrogen gas survive prolonged periods of nitrogen starvation in a chlorotic, dormant state where cell growth and metabolism are arrested. Upon nutrient availability, these dormant cells return to vegetative growth within 2–3 days. This resuscitation process is highly orchestrated and relies on the stepwise re-installation and activation of essential cellular structures and functions. We have been investigating the transition to chlorosis and the return to vegetative growth as a simple model of a cellular developmental process and a fundamental survival strategy in biology. In the present study, we used quantitative proteomics and phosphoproteomics to describe the proteomic landscape of a dormant cyanobacterium and its dynamics during the transition to vegetative growth. We identified intriguing alterations in the set of ribosomal proteins, in RuBisCO components, in the abundance of central regulators and predicted metabolic enzymes. We found O-phosphorylation as an abundant protein modification in the chlorotic state, specifically of metabolic enzymes and proteins involved in photosynthesis. Non-degraded phycobiliproteins were hyperphosphorylated in the chlorotic state. We provide evidence that hyperphosphorylation of the terminal rod linker CpcD increases the lifespan of phycobiliproteins during chlorosis.

## Introduction

Cyanobacteria are among the most widespread prokaryotes on earth, where they inhabit almost the entire illuminated biosphere (1). Their success is based on an autotrophic lifestyle that employs oxygenic photosynthesis and has minimal nutritional requirements. During their evolutionary radiation, a large diversity of cyanobacterial species evolved that are perfectly adapted to their niches. Their morphologies range from simple unicellular to complex filamentous organisms (2).

Cyanobacteria can be divided into two physiological groups with regard to their nitrogen demand, namely diazotrophs and non-diazotrophs. Diazotrophic cyanobacteria can utilize ubiquitous di-nitrogen gas via their nitrogenase activity; this enzyme is protected against photosynthetically produced oxygen through various strategies (3). By contrast, non-diazotrophic cyanobacteria depend on a combined nitrogen source, such as nitrate or ammonium.

In many aquatic environments, nitrogen availability can be highly variable during seasonal cycles. To cope with periods of nitrogen depletion, non-diazotrophic freshwater cyanobacteria, such as *Synechococcus elongatus* PCC 7942, *Synechocystis* sp. PCC 6803 (hereafter *S. elongatus* and *Synechocystis* sp.) or *Arthrospira* sp. PCC 8005 use a special strategy known as chlorosis (4–6). Chlorosis represents a stepwise quiescence mechanism that resembles the widely used prokaryotic survival strategy of the formation of dormant states (7). A hallmark of chlorosis is the rapid degradation of the abundant phycobilisome light-harvesting complexes, initiated by the expression of the gene encoding the Clp-protease adaptor protein NblA (8). This immediate response releases amino acids for transient maintenance of protein synthesis required for stress acclimation. When nitrogen starvation persists, the cells gradually turn down metabolism and enter a dormant-like state characterized by minimum residual photosynthesis and pigmentation, which allows survival of prolonged periods of time (9,10). During that stage, cells maintain a substantial amount of mRNA (11), in spite of highly reduced protein synthesis (9), suggesting that these transcripts are either not translated or translated very slowly.

Following the addition of a nitrogen source, the cells rapidly awaken from dormancy and return to vegetative growth (11). Analysis of transcriptome dynamics during resuscitation, along with physiological and electron microscopy studies have revealed first insights into a genetically determined program guiding resuscitation. After nitrate addition, the cells first activate respiration and catabolize stored glycogen. The first genes to be highly induced (at the transcriptional level) are those for the entire translation apparatus and those for central anabolic processes. Following this initial phase, which lasts about 12–16 h, a metabolic re-wiring for the photosynthetic lifestyle occurs, as evidenced at the transcriptome level and manifested by the respective cellular activities. At this point, the cells start to re-green as the photosynthetic apparatus reconstitutes and finally resume growth, 48–72 h after nitrate addition. Almost all cells in a population escape from chlorosis together. Therefore, the transition from and to chlorosis can be regarded as a reversible developmental process.

A widespread regulatory mechanism that directs developmental processes in Eukarya is protein O-phosphorylation (on Ser/Thr/Tyr residues). It was long believed that in Bacteria, by contrast, only other forms of phosphorylation, such as that of two-component systems (on Asp/His residues) were relevant. In cyanobacteria, several phosphoproteins, mainly involved in photosynthesis, had been reported (12–14). In 1994, the first site-specific bacterial O-phosphorylation event was described in these bacteria, namely on seryl residue 49 of the nitrogen regulatory P_II_ protein (15). With recent developments in mass spectrometry (MS) based phosphoproteomics, it became clear that O-phosphorylation was widely distributed also in the domain Bacteria (16,17). The first qualitative phosphoproteome studies of unicellular cyanobacteria reported large datasets of O-phosphorylation events (18,19). However, such studies are limited to snapshots of the phosphoproteome under a specific condition. To investigate the dynamic changes in O-phosphorylation events in response to defined stimuli, quantitative phosphoproteomic studies are required. Indeed, in the first quantitative study of *Synechocystis* sp., wide-ranging dynamic changes of site-specific phosphorylation levels in response to nitrogen deprivation could be detected (20). We therefore suspected that O-phosphorylation plays an important role also during the resuscitation from chlorosis.

The resuscitation of chlorotic cyanobacteria reveals interesting parallels to the germination of plant seeds, such as the early onset of metabolism. A key feature in seed germination is the selective translation of stored mRNAs (21,22). To solve if selective translation of stored mRNAs is also relevant in the awakening of chlorotic cyanobacteria, knowledge of the proteome dynamics is required. In this study, we addressed this issue by performing quantitative (phospho)proteomics to elucidate global proteome and phosphorylation dynamics during the resuscitation process. Resting on the groundwork of our previous quantitative study, proteome dynamics shed new light on the reconstitution of cellular functions. Our results indicated that protein O-phosphorylation plays an important role in the developmental program and affects the persistence of residual light-harvesting structures during chlorosis.

## Experimental Procedures

### *Synechocystis* sp. Cultivation and Sampling

Wild-type *Synechocystis* sp. PCC 6803 was propagated photoautotrophically under constant illumination with 40 μmol photons m^−2^ s^−1^ at 26 °C in BG11 medium (23) supplemented with 5 mM NaHCO_3_ and 5 mM HEPES without additional CO_2_ supplementation. These growth conditions result in typical linear growth curves of light-limited cultures with initial generation times of about 16 h. Batch cultures of 2.5 L were shaken or stirred at 120 rpm and bubbled with ambient air. Evaporation was compensated for by adding ddH_2_O. At an optical density (OD_750_) of 0.6, nitrogen chlorosis was induced by shifting to nitrogen-free medium as described elsewhere (20). Resuscitation of dormant cultures was triggered by addition of NaNO_3_ (final concentration 10 mM). Cells were harvested at indicated time points as described previously (20). In brief, culture volumes of 400 (T8, T24) or 500 mL (T0, T2, T55) were sampled and rapidly cooled on ice to 0 °C. Cells were then pelleted by centrifugation at 7,477 ×*g* for 8 min. The supernatant was removed, and cells were washed with ice-cold PBS buffer. Cell pellets were snap frozen in liquid nitrogen and stored at −80 °C.

### Construction of *Synechocystis* sp. *cpcD* Point Mutants

The genomic *cpcD* gene in wild-type *Synechocystis* sp. was replaced by homologous recombination with mutant genes encoding either alanine (CpcD^Ala^) or aspartate (CpcD^Asp^) in place of both seryl residues at positions 5 and 11 as follows. The mutant *cpcD* genes were constructed from four overlapping DNA fragments (Table 1), which were amplified from *Synechocystis* sp. genomic DNA as template and ligated by Gibson assembly (24). Each *cpcD* construct, with homologous upstream and downstream regions (~500 bp) and a spectinomycin resistance cassette (Spec^R^), was inserted in the XbaI site of plasmid pUC19. The resulting plasmids, pUC19-cpcD^Ala^ and pUC19-cpcD^Asp^, were transferred by electroporation into *Escherichia coli* DH10B™ cells (ThermoFisher), which were then cultivated in the presence of ampicillin and spectinomycin (each 50 μg mL^−1^). Plasmid DNA was isolated, purified, and transferred into *Synechocystis* sp. wild-type cells via their natural competence. Mutant cells were selected and allowed to segregate on BG11 agar plates with increasing spectinomycin concentrations from 10 to 150 μg mL^−1^. Segregation of mutants was validated by sequencing the PCR-amplified *cpcD* gene fragment from isolated genomic DNA (using primers CpcD^Seq^-F: ATTCATACGGGCATAGGG and CpcD^Seq^-R GAGGAGAAGTGGCTTGAC). Stock cultures of the mutants were kept in BG11 medium with 50 μg mL^−1^ spectinomycin. Experimental cultures were propagated and starved in medium without antibiotics. Mutant and control (wild-type) cells for proteomic experiments were sampled during vegetative growth (OD_750_ = 0.6), during chlorosis (7 d nitrogen depletion), and during resuscitation (48 h after adding nitrate).

**Table 1.**
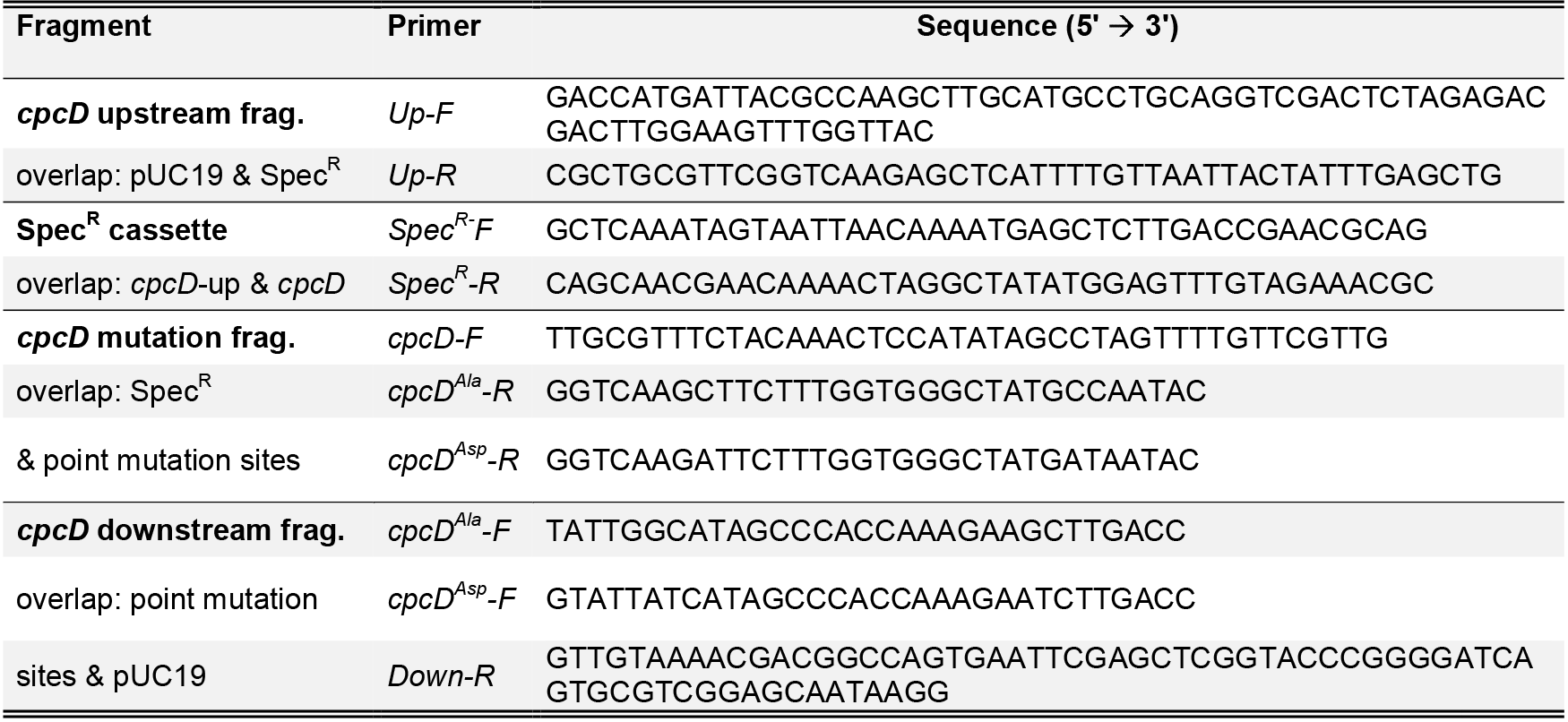
DNA oligonucleotides used for construction of cpcD variants.

### Absorption and 77K Fluorescence Measurements

Optical densities and absorption spectra (400–750 nm; adjusted to OD_750_ = 0.4) of *Synechocystis* sp. cultures were recorded with a spectral resolution of 1 nm on a Speccord^®^ 205 UV/VIS spectrophotometer (Analytic Jena). For 77K fluorescence measurements, cells from 250 mL batch cultures were sampled at the indicated time points and adjusted to an OD_750_ of 1 by dilution with nitrogen-free BG11 medium or concentration by centrifugation at 3,500 ×*g* for 10 min and resuspension in nitrogen-free BG11 medium. Aliquots of 5 mL were transferred to glass tubes (6 mm inner diameter), snap frozen in liquid nitrogen, and stored at −80 °C. Fluorescence spectra were recorded on an AMICO Bowman^®^ Series2 Luminescence Spectrometer at 77K. Fluorescence excitation of Chl *a* and phycobilisomes was at 440 and 580 nm, respectively, and fluorescence emission was recorded at 630–780 nm with a 4 nm detection bandwidth and 1 nm s^−1^ detection speed. Fluorescence detector sensitivity was set to 60% of full scale. Spectra of each sample were measured three times and integrated by the device-specific AB2 software. The detector voltage was set to 880 volts at pigment absorption peaks of 725 nm (Chl a) or 580 nm (phycobilisome). For normalization, the fluorescence intensity of fresh BG11 medium was measured and subtracted from acquired curves, followed by spectral smoothing (based on averaging each value with the three following values).

### Pulsed Amplitude Modulation (PAM) Fluorometry

PAM fluorescence of the minimum PSII fluorescence F_0_ and the PSII quantum yield in light-curve experiments were measured at a Water-PAM chlorophyll fluorometer (Walz GmbH), as described previously for *Synechocystis* sp. (25).

### Experimental Design and Statistical Rationale

For analysis of phosphoproteome dynamics of wild-type cells during resuscitation, two independent biological replicates were analyzed by mass spectrometry (see Figure 2) and raw data was processed by MaxQuant software as described below. Proteome data comprised a total of 36 raw files (18 from each replicate), and phosphoproteome data a total of 40 raw files (20 from each replicate). For comparison in the abundance of photosynthetic proteins between CpcD mutants and wild-type cells, two independent biological replicates were analyzed by mass spectrometry for chlorotic and resuscitation conditions. At vegetative conditions, where the phycobiliproteins were not divergent between the wild-type and mutants, one quantitative measurement was analyzed.

### Sample Preparation for Quantitative Phosphoproteomics

Proteins of each sample were extracted in SDS buffer, precipitated, and pre-digested with Lys-C for 3 h prior to trypsin digestion o.n. as described previously in Method B (20). The resulting peptide solutions were acidified with trifluoroacetic acid (TFA) to pH 2.5 and loaded onto SepPak C_18_ cartridges (Waters) for desalting and subsequent dimethylation labeling (26). Labeling efficiencies were validated (**Supplementary Figure 1**) prior to the mixing of differentially labeled samples in a 1:1:1 ratio (light: intermediate: heavy label), based on quantified protein amounts from Bradford assay and pilot MS measurements. For phosphoproteome analysis, a total of 5 or 12 mg peptides, originating from 2.5 or 5 L culture, respectively, was utilized per experiment (for the first and second replicate, respectively). For proteome measurements, each 50 μg of the peptide mixtures was fractionated by reversed-phase chromatography at high pH (Pierce™ High-pH Reversed-Phase Peptide Fractionation Kit), according to the manufacturer’s protocol, except that peptides were eluted in nine fractions of acetonitrile in 10 mM ammonia instead of triethylamine (5, 7.5, 10, 12.5, 13.3, 15, 17.5, 20, and 50%, v/v). Peptide fractions were purified on C_18_ stage tips (27) and then analyzed by nanoLC-MS/MS. For quantitative comparison of photosynthetic proteins of CpcD mutants and wild-type, 0.1 mg protein per sample was labeled and analyzed likewise. Per condition, two independent samples were measured, and protein ratios were averaged due to similar trends. For phosphoproteome measurements, phosphopeptides were enriched by TiO_2_ chromatography in ten enrichment cycles as described previously (20). In brief, peptide mixtures in 80% (v/v) acetonitrile (aq.) were supplemented with TFA to a final concentration of 6% (v/v), and incubated for 10 min with TiO_2_ spheres (Sachtopore NP 5 jm diameter; protein-to-TiO_2_ sphere ratio of 10:1) on an orbital shaker. Spheres were subsequently washed with 200 μL 80% acetonitrile/6% TFA (v/v) solution twice and loaded onto C_8_ (Empore™) stage tips. After flushing with 100 μL 80% acetonitrile/1% TFA (v/v), phosphopeptides were eluted in three steps: with 30 μL of 1.25% and then 70 μL of 5% (v/v) ammonia solution (aq.), and 20 μL 60% acetonitrile/1% TFA (v/v). Eluted phosphopeptides were mixed with 30 μL 20% TFA (aq.). Eluates were purified on C_18_ stage tips and analyzed by nanoLC-MS/MS.

### Mass Spectrometry Measurements

For proteome measurements, fractionated samples were loaded onto an RP C_18_ nanoHPLC column (20 cm, 75 μm inner diameter PicoTip fused silica emitter (New Objective) in-house packed with 1.9 μm ReproSil-Pur C18-AQ resin; Dr. Maisch) on an EASY-nLC 1200 system (Thermo Scientific). Peptides were separated by fraction-specific 90-min segmented linear gradients (optimized for increasing sample hydrophobicity; **Supplementary Figure 2**). Eluted peptides were ionized and analyzed on an electrospray ionization (ESI) source coupled on-line to a Q Exactive HF mass spectrometer (ThermoFisher). Spectra were acquired in the positive-ion mode with a mass range *m/z* of 300–1,650 at a resolution of 60,000. The 12 most intense multiple charged ions were selected for fragmentation by higher-energy collisional dissociation (HCD), and MS^2^ spectra were recorded at a resolution of 30,000. For phosphoproteomes, the first replicate was measured on an Orbitrap Elite mass spectrometer and the second replicate was measured on a Q Exactive HF mass spectrometer (both ThermoFisher). The ESI source of the Orbitrap Elite was coupled to an EASY-nLC II system (Proxeon Biosystems), on which enriched phosphopeptides were loaded onto a reversed-phase C_18_ nanoHPLC column (15 cm column in-house packed with 3 μm ReproSil-Pur C18-AQ resin). Phosphopeptides were eluted using a segmented 87-min gradient of 5–33% of HPLC solvent B (80% acetonitrile in 0.5% formic acid) at a constant flow rate of 200 nL min^−1^. MS scans were acquired in the positive-ion mode at *m/z* 300–2,000 at a resolution of 120,000. The 15 most intense multiple-charged ions were selected for fragmentation by HCD, and MS^2^ spectra were recorded at a resolution of 15,000. The Q Exactive HF-ESI source was coupled to an EASY-nLC 1200 system; phosphopeptides were loaded and eluted as described above. MS full scans were acquired at *m/z* 300–1,650 and a resolution of 60,000. Ions were selected and fragmented for MS^2^ spectra as described above at a resolution of 60,000, and the maximum ion injection time was set to 220 ms. Dynamic exclusion of sequenced precursor ions for 90 s (Orbitrap Elite) or 30 s (Q Exactive HF) was enabled.

### Data Processing and Validation

Raw MS spectra were processed with MaxQuant software suite (version 1.5.0.12) (28) at default settings. Identified peaks were searched against a target-decoy database of *Synechocystis* sp. PCC 6803 with 3,671 protein sequences, retrieved from Cyanobase (http://genome.microbedb.jp/cyanobase) (29) (version 06/2014), and against 245 common contaminants. The following database search criteria were defined. Trypsin was defined as a cleaving enzyme, and up to two missed cleavages were allowed. Dimethylation on peptide N-termini and lysine residues was defined as light (+28.03 Da), intermediate (+32.06 Da), and heavy (+36.08 Da) labels. Carbamido-methylation of cysteines was set as fixed modifications, and methionine oxidation, protein N-termini acetylation, and Ser/Thr/Tyr phosphorylation (only in phosphoproteome analyses) were set as variable modifications. Re-quantification and match between runs options were enabled, and the initial mass tolerance of precursor ions was limited to 20 ppm and 0.5 ppm for fragment ions. False discovery rates retrieved from MaxQuant of peptides and proteins were each limited to 1%, and quantitation of labeled peptides required at least two ratio counts (**Supplementary peptides table**). Peptides and phosphopeptides were only allowed with posterior error probabilities (PEP) <1%. Proteins identified by a single peptide are indicated in the dataset and MS^2^ spectra are provided (**Supplementary MS^2^ spectra of proteins identified based on a single peptide**). Phosphopeptide MS^2^ spectra were manually filtered and validated with stringent acceptance criteria, namely both a comprehensive coverage of b-and y-ion series for HCD spectra and a low noise-to-signal ratio were required (**Supplementary phosphopeptide MS^2^ spectra**). Furthermore, the localization of the phosphorylation site was validated. To exclude quantitation bias of phosphopeptide ratios due to fluctuations of corresponding proteins, phosphopeptide ratios were normalized by dividing the corresponding protein ratios of each experiment. Protein levels were analyzed by Pearson correlation with Perseus software (version 1.5.0.31).

### Cluster Analysis

Similar protein and phosphorylation dynamics were grouped in profiles using a hierarchical clustering approach. The standard deviations of log2-transformed protein and protein O-phosphorylation event (p-event) data, quantified at all five time points in the regeneration process, were calculated as a measure of their abundance change. For the proteome, only proteins among the top 25% of calculated standard deviations were considered to ensure a substantial abundance change. Cluster analysis was performed with the “hclust” R function on Z-score-transformed profiles using Euclidian distance as metric in combination with the “ward” clustering method as described previously (30). For the analysis of protein functions, available annotations of the *Synechocystis* sp. proteome were retrieved via the Perseus software suite. Each cluster was manually functionally assigned based on all available annotations.

## Results

To gain deeper insights into the developmental process of chlorosis and resuscitation of cyanobacteria, we aimed at a quantitative analysis of the (phospho)proteome in the course of resuscitation form chlorosis. As larger sample volumes are required for this analysis than for our previous studies (5,9,11), the appropriate sampling points had to be identified under non-stressed, light-limited growth conditions. *Synechocystis* sp. cultures were shifted to medium lacking combined nitrogen, and the progress of chlorosis was followed until a stable residual level of pigmentation and metabolic quiescence was reached after 21 days. At that time point (T0), resuscitation was started by addition of nitrate. We monitored the cell density and re-pigmentation of the cultures until full recovery (**Supplementary Figure 3A**). Photosynthesis re-initiated after 24 h (T24), and the cell density began in increase after 48 h (T48) (**Supplementary Figure 3B**). To address the timing of the earliest cellular activities, we determined the utilization of supplemented nitrate. When chlorotic cells start to reduce nitrate, traces of nitrite in the medium can be measured (31). Remarkably, elevated nitrite concentrations were detected already 15 min after nitrate addition, which demonstrated rapid nitrate conversion and an almost immediate physiological response of dormant cells to a nitrogen source (**Supplementary Figure 4**).

Whole-cell absorbance spectra over the course of resuscitation indicated that phycobiliproteins increased from an initial very low basal level to a 25-fold higher level over the course of resuscitation, and Chl *a* (in photosystems I and II; PSI and PSII) increased from a higher residual level to a 3.5-fold higher level (**Supplementary Figure 5A and B**), similar to our previously reported results (see Introduction and (11)). We used 77K fluorescence spectroscopy to further investigate the regeneration of the photosynthetic apparatus. Specific excitation of Chl *a* at 440 nm or phycobilisomes at 580 nm allows one to assess the assembly and functionality of all components involved in light-energy transduction to the reaction centers. In vegetative cells 72 h after initiation of resuscitation (T72), excitation of Chl *a* at 440 nm (Figure 1A) yielded a dominant fluorescence of PSI Chl *a* (725 nm) emission peak and minor emission of PSII core antennae at 685 and 695 nm, as described for intact PSII complexes (32). In the chlorotic state, substantial PSI fluorescence was detected, whereas PSII fluorescence was reduced to almost undetectable levels. This relatively high PSI signal agrees with the residual 30% Chl *a* in chlorotic cells because approximately 85% of Chl *a* is associated with PSI, but only a small fraction is associated with PSII (33). Consequently, the increase in Chl *a* during resuscitation (Supplementary Figure 5B) parallels the increase in PSI fluorescence.

**Figure 1:**
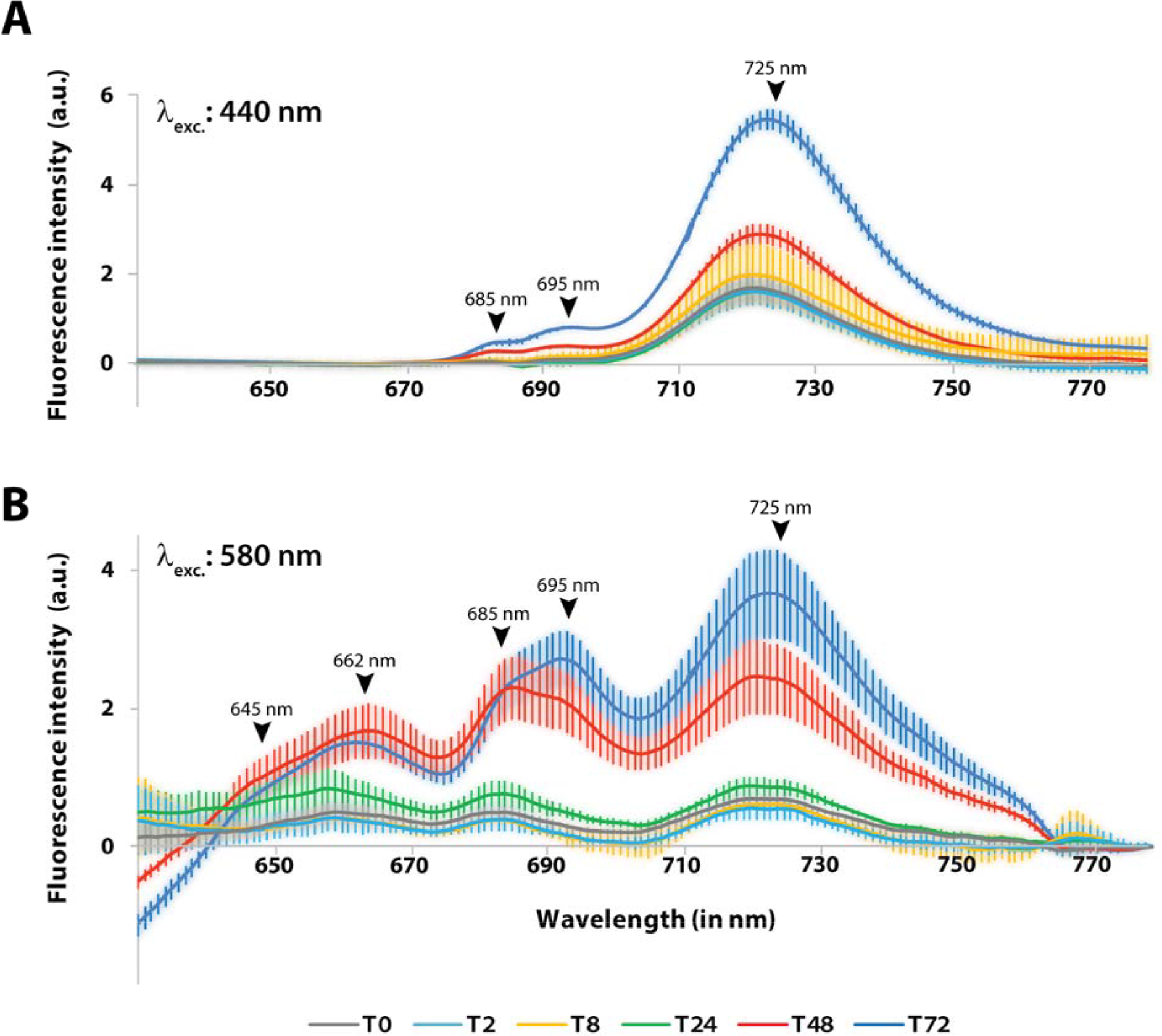
77K fluorescence spectroscopy of *Synechocystis* sp. cultures during the regeneration process. Fluorescence emission of samples was recorded at the indicated time points at 77K after excitation of **(A)** Chl *a* in PSI at 440 nm or **(B)** phycobilisomes at 580 nm. The fluorescence emission spectra shown are the average of three independent replicates, each of which was measured three times. Error bars indicate the standard error of mean. Peaks of fluorescence signals of photosynthetic pigments are indicated: phycocyanin, 645 nm; allophycocyanin, 662 nm; PSII core antennae, 685 nm and 695 nm; PSI Chl a, 725 nm.

When phycobiliproteins are excited at 580 nm (Figure 1B), fluorescence is emitted from phycocyanin and allophycocyanin, PSII core antennae (CP43, CP47), and PSI Chl *a*. The corresponding signals were clearly visible in recovered cells 72 h after initiation of resuscitation. By contrast, chlorotic cells displayed very low, but clearly detectable fluorescence emission, corresponding to the strongly reduced phycobiliprotein content. This congruence indicates that the remaining phycobiliproteins still function in light harvesting and transferring of energy to remaining PSII and PSI. The first increase in PSII and PSI fluorescence emission became visible after 24 h of resuscitation. After 48 h of resuscitation, fluorescence emission almost reached that of fully restored cells, but the ratio between emission at 685 nm and 695 nm (originating from CP43 and CP47; (32)) was shifted, which indicated that PSII complexes were still immature (see Discussion and (34)).

### Design of the Quantitative (Phospho)Proteome Experiments

To determine the (phospho)proteome dynamics during resuscitation from long-term chlorosis, we took samples at five time points (**Supplementary Figure 6**): T0, the dormant state before nitrate addition; T2, the early awakening, 2 h after nitrate addition; T8, the glycolytic state; T24, the transition to photosynthesis; and T55, the return to exponential growth. Our quantitative proteomic workflow included dimethylation labeling, which allowed quantitative comparison of three samples in one analysis (26). The samples of the five time points were assigned to two separate triplex-labeling experiments (T0-T24-T55 and T2-T8-T55), with sample T55 used as a common reference point (Figure 2). The ratios between the three labelings (light, intermediate, and heavy) were consequently normalized to the reference point (defined as 1). The analysis of two biological replicates resulted in a combined dataset, comprising 2,461 proteins overall (False discovery rate of 0.28% at the peptide level and 1.44% at the protein level; **Supplementary Table S1**). The proteome coverage corresponds to 67% of the theoretical *Synechocystis* sp. proteome (29). Hereof, 2,271 proteins yielded quantitative data (including all five time points). The two replicates were reproducible, as shown by paired correlation analysis, which indicated that protein dynamics were consistent (**Supplementary Figure 7**) and that the quality of the data corresponds to the standards of stable isotope labeling/MS studies. Calculated Pearson coefficients were generally higher between earlier than between later time points, which indicates that dormant cells contain a defined proteome and that the initial phase of resuscitation proceeds without any delay between replicates. Increasing temporal variations towards the second phase of the resuscitation process (at T24) are indicated by declining correlation coefficients between replicates.

**Figure 2:**
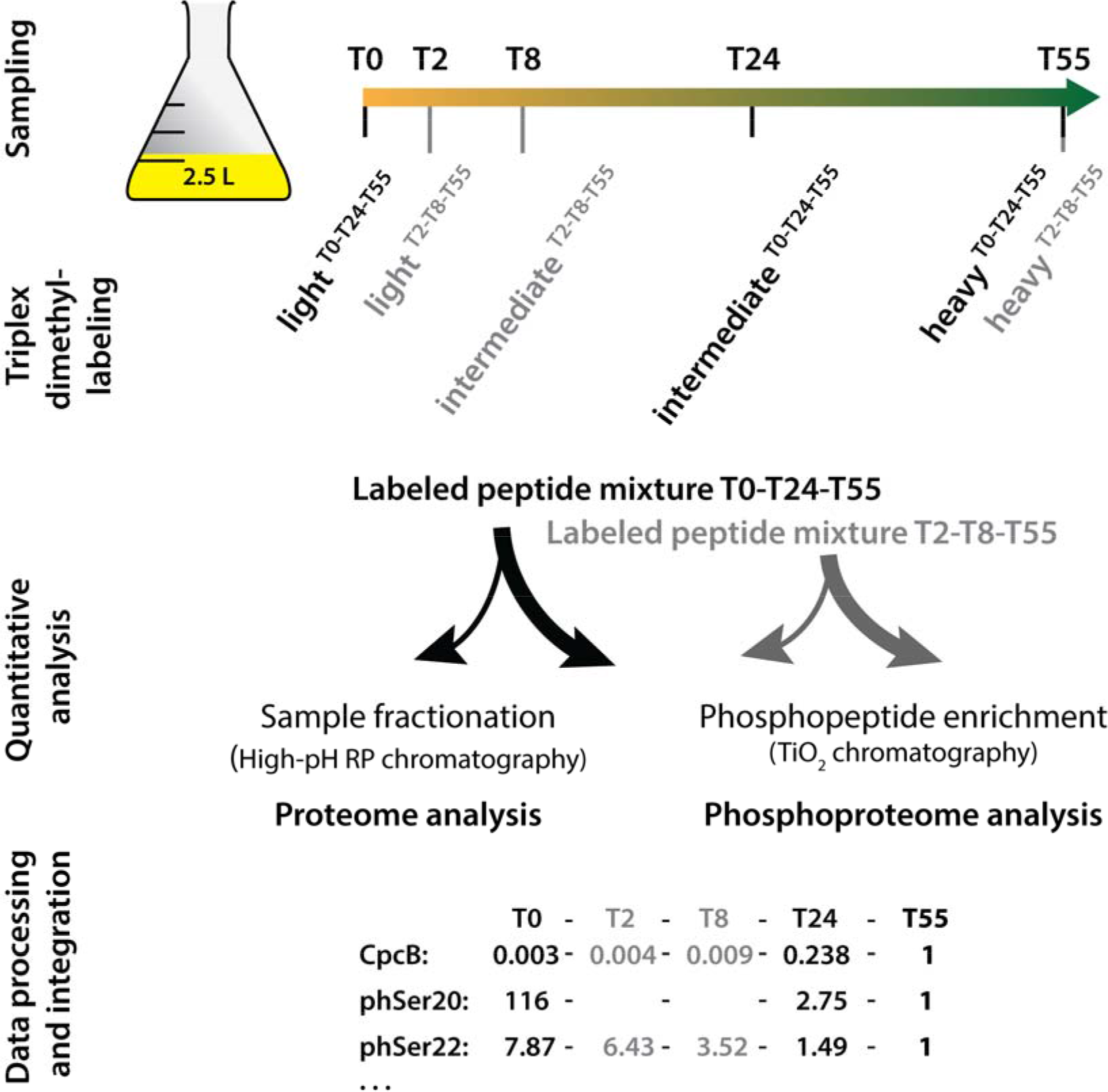
Schematic representation of the phosphoproteomic study design. All five time-point samples of one replicate originate from the same culture; growth curves of replicate cultures are shown in Supplementary Figure 4. Samples from time points T0-T24-T55 (indicated in black) and T2-T8-T55 (indicated in grey) were separately triplex labeled, and the corresponding light, intermediate, and heavy dimethyl labels are indicated. Labeled peptide mixtures were combined in equal amounts per triplex mix. Peptides in an aliquot of each mixture were fractionated for consecutive proteome analysis, and the remainder was used to enrich phosphopeptides for phosphoproteome analysis. Mass spectroscopy data on relative protein and O-phosphorylation abundances relative to those of the reference point T55 were integrated into the full dataset, as indicated exemplarily for the CpcB protein and two phosphorylation events on Ser20 and Ser22.

### The Proteome of Long-term Starved Chlorotic Cells

In the quantitative proteome dataset (Supplementary Table S1) the ratio compares the abundance of a particular protein at any sampling point relative to T55. The total amount of protein isolated from cultures of long-term chlorotic cells (T0) was about fourfold lower than in T55 cells (normalized to OD_750_) (**Supplementary Figure 8**). For quantitative analysis, the samples must be adjusted and normalized to the same amount of protein, which leads in this case to an overrepresentation of the chlorotic cells. As expected, the proteins with the strongest reduction during the chlorotic state were phycobiliproteins. Of these, the strongest decrease was detected for the peripheral phycobilisome rods (T0:T55 = 0.003–0.08), which represent the five most strongly diminished proteins of the entire proteome during chlorosis. The phycobilisome core components were about three-to fourfold higher in abundance (T0:T55 = 0.011–0.028). Only the phycobilisome core membrane linker ApcE remained at slightly higher levels (T0:T55 = 0.47). These differences indicate that the rod/core ratio of the residual phycobilisome is about three-to fourfold lower than in phycobilisomes in cells grown under standard conditions, which is in agreement with a classical study on isolated phycobilisomes (35).

Proteins belonging to PSII were in general more diminished during chlorosis than those of PSI. The oxygen-evolving-complex protein PsbP showed the strongest reduction, (T0:T55 = 0.14), and other essential PSII proteins were diminished with ratios of 0.199–0.684. In agreement with fluorescence spectra, the amounts of most structural PSI components remained high in the chlorotic state (T0:T55 = 0.9–1.9). Interestingly, the alternative PSI reaction center subunit X protein PsaK2 accumulated in chlorosis (T0:T55 = 1.9), whereas the abundance of standard subunit X was so low that it could not be quantified. PsaK2 is essential for acclimation of PSI to high light levels (36). Similarly, the strong decrease in the accessory PSI subunit IX (T0:T55 = 0.133) indicated a modified PSI complex in chlorotic cells. Altogether, the proteome of the photosynthetic machinery indicated that in chlorotic cells, several properties of the residual photosynthetic apparatus resemble those of high-light-stressed cells (phycobilisome rod reduction, PSI alteration).

Other known proteins whose amounts were strongly decreased in chlorotic cells comprised, among others, the glutamine-synthetase-inactivating factor 7 (IF7), cytochrome *c*_553_, and oxygen-dependent coproporphyrinogen III oxidase (HemF). A decrease in IF7 leads to high glutamine synthetase activity; low levels of cytochrome *c*553 correlates with reduced photosynthetic electron transport, and depression of HemF reflects a reduced rate of tetrapyrrole biosynthesis. Furthermore, several key metabolic enzymes were strongly reduced in the chlorotic state, such as the acetyl-CoA carboxylase alpha and beta subunits, pentose-5-phosphate-3 epimerase, and S-adenosylmethionine synthetase, all of which displayed T0:T55 ratios of 0.09 or less. This decrease correlates well with an arrested anabolism. The low level of particularly these enzymes might indicate a regulatory hot spot at these metabolic sites. Intriguingly, histidine kinase Sll0750, also termed SasA, was the most strongly depressed two-component regulatory protein during chlorosis (T0:T55 = 0.08). SasA interacts with the circadian clock protein KaiC, thereby linking gene expression to the circadian cycle. Its extremely low abundance in chlorotic cells could indicate that in the chlorotic state, the low level of residual gene expression is not controlled by the circadian clock.

Our previous transcriptome analysis (11) revealed a strongly reduced expression of genes coding for components of the translation apparatus. Although the present proteome analysis confirms the general trend, it provides a more complete picture. In contrast to the uniformly reduced abundances of phycobiliproteins in the chlorotic state, the abundance of ribosomal proteins was divergent, ranging from T0:T55 ratios of 0.11 (L9) to 1.31 (S12), which indicates a differential turnover of the various ribosomal proteins (for a more detailed reflection on ribosomal proteins, see Discussion).

On the other end of the protein abundance range, we found proteins that were markedly enriched in the chlorotic state. Thirty-seven proteins displayed T0:T55 ratios >four. Twenty-five of these (68%) are annotated as “hypothetical” or “unknown”. Of the ten most highly enriched proteins in the chlorotic state, eight are unknown or hypothetical proteins, one has a predicted function as an ATP-binding protein of an ABC transporter (Sll1001), and one is the type 2 NAD(P)H dehydrogenase Slr0851. The strong enrichment of these proteins suggests that they might have important functions in the chlorotic state.

### Proteome Dynamics During Resuscitation

The abundances of most proteins showed a general trend, namely that their abundances did not change much during the first 8 h of resuscitation, but generally increased from T24 to T55. To characterize the dynamic rearrangement of the proteome during the resuscitation process in more detail, we cluster analyzed the proteins with the top 25% ratio changes (based on the dynamic change during the resuscitation, see Methods). This procedure resulted in the unbiased classification of 568 proteins in six defined clusters (**Supplementary Tables S2-S7**), for which protein functions were assigned (Figure 3). Proteins in cluster 1 accumulated early, peaking at T24. In agreement with the early expression of the corresponding genes (11), this cluster is dominated by components of the translational machinery, especially ribosomal proteins, which fulfills the basic requirement for increasing *de novo* protein synthesis. An important aspect of unbiased cluster analysis of expression profiles in large datasets is the possibility to assign potential functions to so far uncharacterized proteins, that are appearing in the same cluster. A dynamic expression profile of a protein of unknown function similar to that of protein of a certain protein complex is an indicator of a functional association. Accordingly, we compared the expression profiles of hypothetical proteins in cluster 1 with ribosomal components. In this way, we identified the Ycf65-like protein (Slr0923), whose transcription and translation profile matches with those of ribosomal components (**Supplementary Figure 9**). Furthermore, many phycobilisome components and anabolic enzymes (nitrogen assimilation, amino acid and fatty acid synthesis) appeared in cluster 1. Proteins in cluster 2 steadily accumulated until T55 at the end of the resuscitation process. This cluster contains many enzymes for pigment synthesis and anabolic enzymes. Proteins in cluster 3, including photosystem and ATPase components, accumulated after some delay, i.e., not increasing before T24, concomitant with the re-appearance of the thylakoid membranes. The appearance of these proteins represents the transition to photosynthetic activity (11). Intriguingly, the few proteins of cluster 4 rapidly but transiently increased, with an almost immediate increase in protein abundance 2 h after initiation of resuscitation. Although most proteins in this cluster are hypothetical proteins, three two-component system histidine kinases were identified, namely Hik8, Hik11, and Hik33, which indicates that these kinases are required early at the start of the resuscitation process.

**Figure 3:**
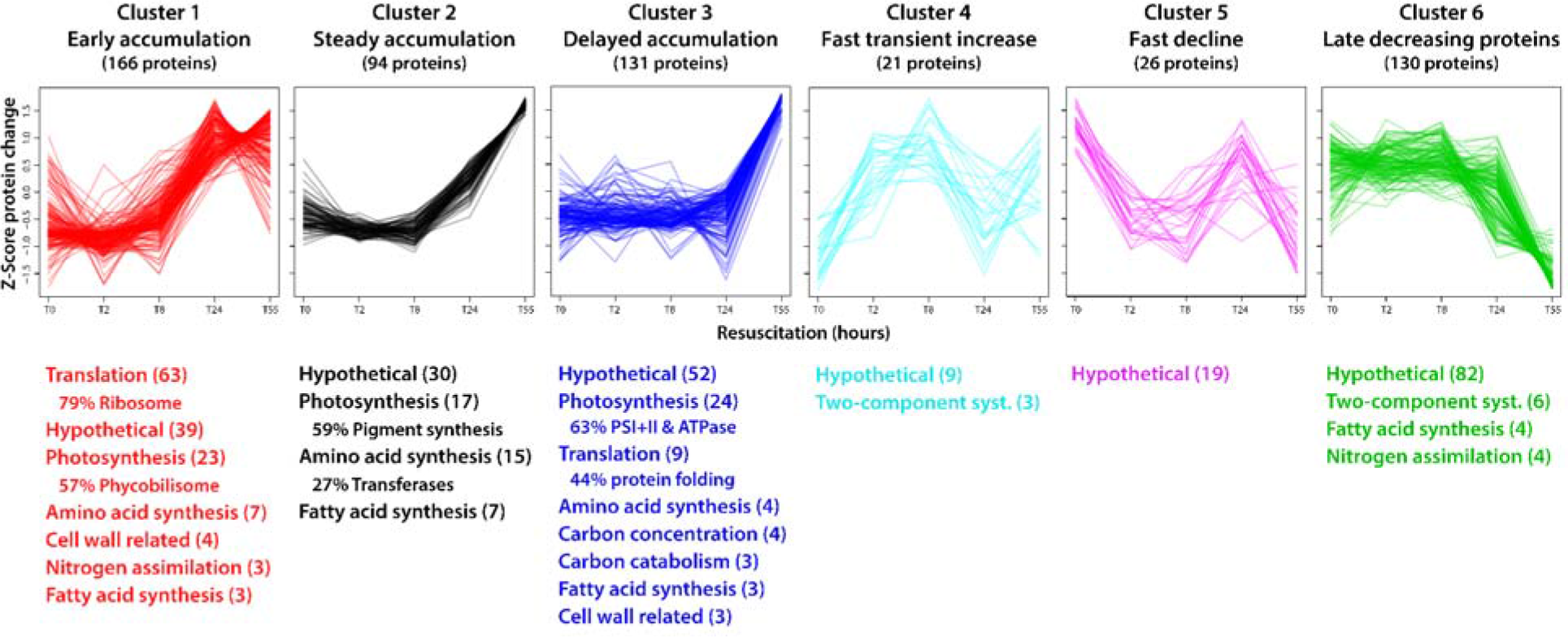
Protein cluster analysis and functional assignment. The proteins with the top 25% changes in the dataset (based on standard deviations of values at five time points) were grouped in six distinct clusters based on profile similarity during the resuscitation process. Assigned protein functions were categorized according to biological processes and are listed below the profiles (with ≥ 3 proteins per category; number of categorized proteins is indicated in parentheses). Where appropriate, the dominant subcategory is indicated in percent relative to the main category. Detailed information of clustered proteins is provided in Supplementary Tables S2-S7.

In contrast to the general trend, abundances of some proteins decreased during resuscitation. These proteins had accumulated during chlorosis, and most are uncharacterized or hypothetical proteins. These proteins were grouped into two distinct clusters: cluster 5 consists of a small group of mainly uncharacterized proteins whose abundances rapidly declined, and cluster 6 consists of a large number of proteins, including several proteins of two-component systems, whose abundances decreased only from T24 to T55. In general, the overall large number of uncharacterized hypothetical proteins in clusters 4–6 underlines our general lack of knowledge of proteins that presumably have important functions in the acclimation to and escape from long-term starvation.

### The Phosphoproteome of Chlorotic Cells

For the analysis of phosphoproteome dynamics, we enriched phosphopeptides by TiO_2_ chromatography and analyzed them by mass spectrometry (Figure 2). We manually validated the phosphopeptide spectra and then identified 186 high-confidence protein O-phosphorylation events (p-events) located on 96 phosphoproteins (**Supplementary Table S8** and Supplementary MS^2^ spectra). Of these, 126 p-events provided quantitative information. We then analyzed the phosphorylation dynamics of the abundant PII signaling protein to validate phosphorylation congruence between replicates. The stepwise phosphorylation of this protein trimer at Ser49 (on three identical subunits) upon nitrogen deficiency serves as a marker for cellular nitrogen availability (15). Using mass spectrometry and non-denaturing PAGE-immunoblot analyses, we observed the expected transition of high to low PII phosphorylation levels throughout resuscitation (**Supplementary Figure 10**). We also identified accessory P_II_-phosphorylation on residues Thr52 and Ser94 by mass spectrometry, but the intensity of these p-events was three orders of magnitude lower than the intensity of Ser49 phosphorylation, and these accessory p-events were more abundant under conditions of long-term starvation. We therefore conclude that these p-events are occasional and not involved in the primary mechanism of P_II_ signal transduction.

Among the top 10 p-events with the strongest enrichment during the chlorotic state, 9 p-events reside in phycobilisome proteins CpcA, CpcB, ApcB, and CpcD (small rod linker), which highlights the hyperphosphorylation of the phycobilisomes remaining during chlorosis (see also below). We detected another highly enriched p-event on phosphoglucomutase (Sll0726), namely a 13.6-fold increase at residue Ser63; this protein also had a highly decreased p-event at Ser168 (T0:T55 = 0.075). Phosphoglucomutase might be a key control point in glycogen catabolism. Other phosphoproteins involved in carbon catabolism that had differential phosphorylation levels are 6-phospho-D-gluconate dehydrogenase (Sll0329; T0:T55 = 1.8), fructose bisphosphate aldolase (Sll0018; T0:T55 = 1.3), and phosphoglycerate mutase (Slr1945; T0:T55 = 0.243). Furthermore, two proteins involved in CO_2_ fixation also showed strongly increased phosphorylation. The CP12 polypeptide, a regulator of the Calvin-Benson cycle, showed increased phosphorylation at two sites; the carboxysome protein CcmM displayed increased phosphorylation at several sites.

Another important functional category of proteins with differential phosphorylation is the ribosomal proteins. The small ribosomal subunit components S5 and S30 showed decreased phosphorylation, whereas the large subunit proteins L9 and L12 displayed p-events with increased abundance.

### O-Phosphorylation Dynamics During Resuscitation from Chlorosis

The unbiased cluster analysis of quantified p-events allowed the assignment of five distinct clusters (clusters A–E; Figure 4; **Supplementary Tables S9-S13**). In most clusters, phosphorylation levels were elevated in the chlorotic state or transiently increased during early resuscitation. By contrast, in the resuscitation phase from T24 to T55, decreasing phosphorylation levels were common for nearly all p-events.

**Figure 4:**
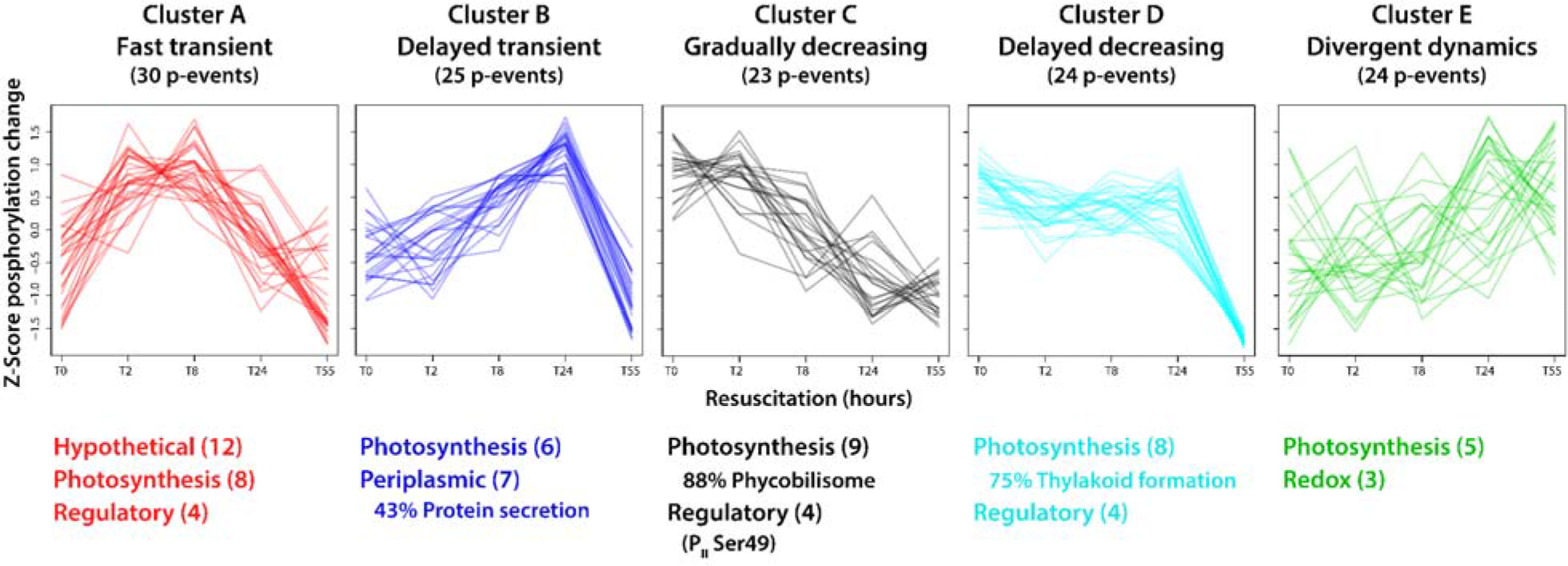
Cluster analysis of phosphorylation events and functional assignment of underlying phosphoproteins. Quantified p-events (n = 126) were grouped into five distinct clusters based on profile similarity during the resuscitation process. Assigned functions of underlying phosphoproteins were categorized according to biological processes and are listed below the profiles (with ≥ 3 p-events per category; numbers of categorized p-events are indicated in parentheses). Where appropriate, the dominant subcategory is indicated in percent relative to the main category. Detailed information of clustered p-events is provided in Supplementary Tables S9-S13.

Cluster A consisted of phosphoproteins with maximal phosphorylation in the early resuscitation phase (T2 and T8); besides a majority of unknown proteins, we find proteins involved photosynthetic processes and with potential regulatory properties (Slr1225, Slr1856, Slr1859, Slr1894). Phosphorylation of proteins of cluster B peaked at T24, and most of these proteins are also involved in photosynthesis or have predicted function in protein secretion. In cluster C, characterized by early dephosphorylation, most phosphorylated phycobiliproteins are present, as well as the nitrogen regulatory PII protein (Ser49 and Thr52) and phosphoglucomutase (Ser63). Cluster D consists of phosphoproteins, which maintain a high level of phosphorylation until T24. Here, we find the Vipp1 thylakoid membrane protein, regulatory proteins (NarL and S/T protein kinase homologues) as well as the CP12 polypeptide. Cluster E is distinct from other profiles and contains proteins with divergent phosphorylation dynamics.

Clearly, components of the photosynthetic apparatus dominated among the detected phosphoproteins. An overview of these phosphoproteins is shown in Figure 5. A total of 17 phosphoproteins are part of the phycobilisome, PSII, and PSI, or are differentially associated with the thylakoid membrane. On these phosphoproteins, altogether 40 p-events were detected, with a high occurrence on phycobiliproteins (19 of 40 p-events), on which multiple phosphorylation sites were frequently detected (e.g., seven on CpcA).

**Figure 5:**
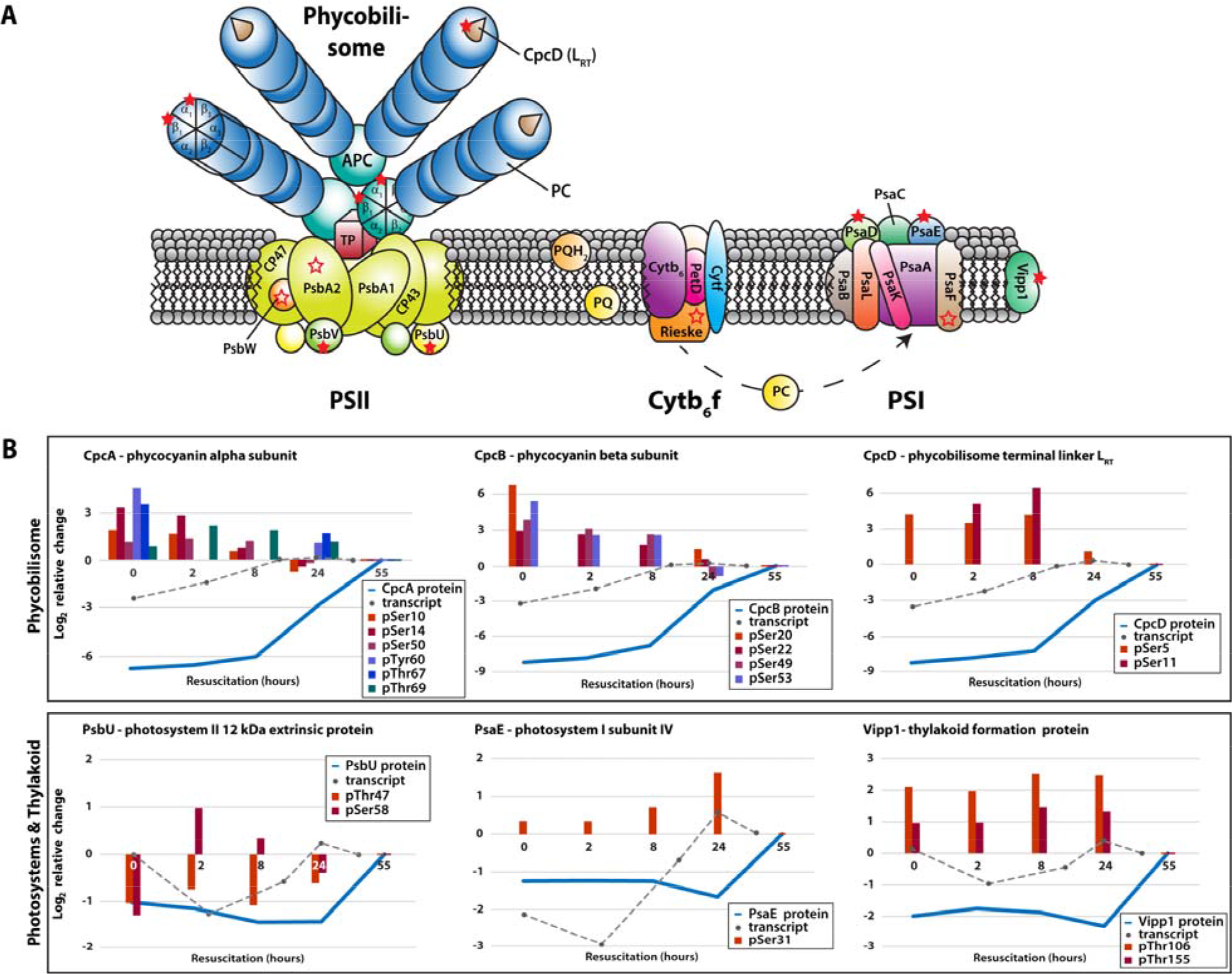
**(A)** Phosphoproteins of the photosynthetic apparatus in *Synechocystis* sp. and **(B)** dynamics of defined components during resuscitation. **(A)** Detected phosphoproteins are indicated by open and filled red stars, including phycobilisomes (PBS), photosystem II (PSII), cytochrome *b_6_f* complex (Cyt *b_6_f*), photosystem I (PSI), and other proteins of the thylakoid membrane. Open red stars indicate phosphoproteins with identified p-events, and filled red stars represent quantified p-events (protein ratios of at least three time points). The photosynthetic apparatus model was adopted from Bryant (83). **(B)** Phosphorylation dynamics of exemplary phosphoproteins during resuscitation on a log_2_xs scale. Blue lines show protein dynamics; gray dotted lines indicate the corresponding transcript dynamics, taken from Klotz et al. (11). Colored bars depict exemplary site-specific phosphorylation dynamics.

As outlined above, the protein levels of phycobilisome components strongly increased during resuscitation, whereas the phosphorylation level decreased. Based on the comprehensive proteome and phosphoproteome data, we calculated phosphorylation site occupancies (Supplementary Table S8) and thereby obtained unprecedented results. Particularly striking were the high phosphorylation site occupancies of the phycocyanin subunits in the long-term chlorotic state, with CpcB being nearly 100% phosphorylated. Rapidly after the onset of resuscitation, phosphorylation site occupancies dropped dramatically and reached minimum levels in the recovered state. We detected similar trends also for multiple p-events on CpcA and for phosphorylation of Ser5 on CpcD (Table 2). Although we could determine the occupancy for only one of two quantified p-events on CpcD, phosphorylation levels at both sites, Ser5 and Ser11, strongly decreased towards the end of resuscitation. Furthermore, a third p-event on Ser36 could be detected, but not quantified.

**Table 2.**
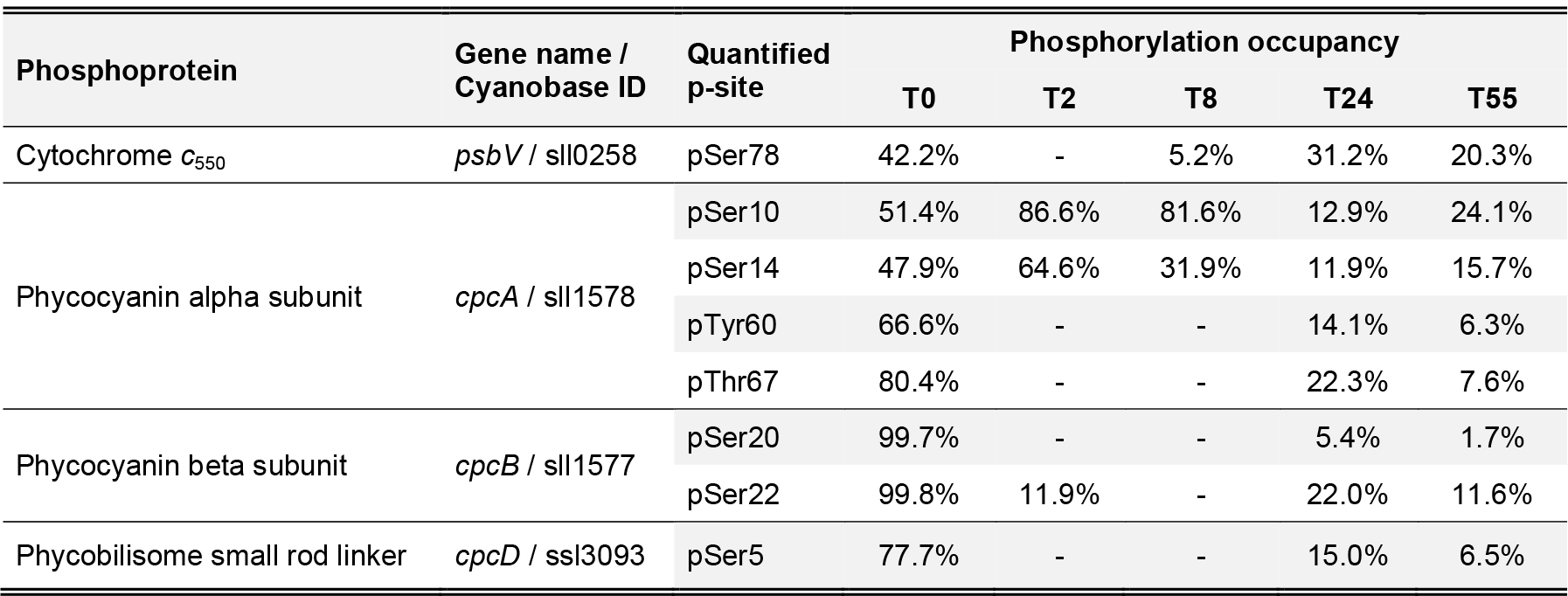
**Photosystem phosphorylation occupancies** (with at least tree measured values)

We observed the most prominent dynamics on both protein and phosphorylation levels during the transition of chlorotic cells to vegetative growth in components of the photosynthetic apparatus. The low abundant residual phycobiliproteins (≤ 1% of phycocyanin and ≤ 3% of allophycocyanin and CpcD, relative to vegetative cells) were in a highly phosphorylated state during chlorosis. These residual phycobilisome complexes remained functional, as revealed by 77K fluorescence spectroscopy. This raised the question of the role of the detected phycobiliprotein hyperphosphorylation for the survival and resuscitation of chlorotic cells.

### Phosphorylation of CpcD Regulates Phycobilisome Turnover

The detected phosphorylation of CpcD, the rod linker that terminates the rods, is in accordance with earlier findings in several cyanobacterial species (18,19,37). We hypothesized that CpcD phosphorylation might play a role in preventing the complete degradation of phycobilisome complexes during chlorosis to maintain a low percentage of functional phycobilisomes. To test this hypothesis, we mutated both seryl residues (Ser5 and Ser11) pairwise to two non-phosphorylatable Ala (CpcD^Ala^) or to Asp (CpcD^Asp^) residues. Although Asp has a lower electrostatic potential than a phosphate group (i.e., one instead of two negative charges), it is commonly used as a potential phospho-mimic.

To study the effects of these CpcD mutations, mutant and wild-type cultures were cultivated under conditions of very low light (5 μmol photons m^−2^ s^−1^) to slow down phycobiliprotein degradation to obtain a better time resolution of the degradation kinetics. As revealed by whole cell absorbance spectra, the CpcD mutations had no effect on phycobiliprotein abundance during vegetative growth (Figure 6A). By contrast, upon nitrogen depletion, phycobiliproteins were degraded faster in both mutants than in the wild-type. At the final chlorotic state after 12 days of starvation, phycobiliproteins were almost undetectable in the mutants, whereas residual amounts were maintained in the wild-type (see also Figure 8). In contrast to the effect of the mutations on phycobiliproteins, the cellular Chl *a* concentrations were slightly higher in the mutants after prolonged chlorosis, whereas they were unaffected during vegetative growth (**Supplementary Figure 11**). Cell viability of the mutants in the chlorotic state was not impaired after 21 days of starvation (**Supplementary Figure 12**), which indicated that the maintenance of residual phycobilisomes during chlorosis is not essential under laboratory conditions. Furthermore, the similar effect of the CpcD^Ala^ and CpcD^Asp^ mutants indicated that the Ser-to-Asp mutation is not a phosphomimetic mutation in this case.

**Figure 6:**
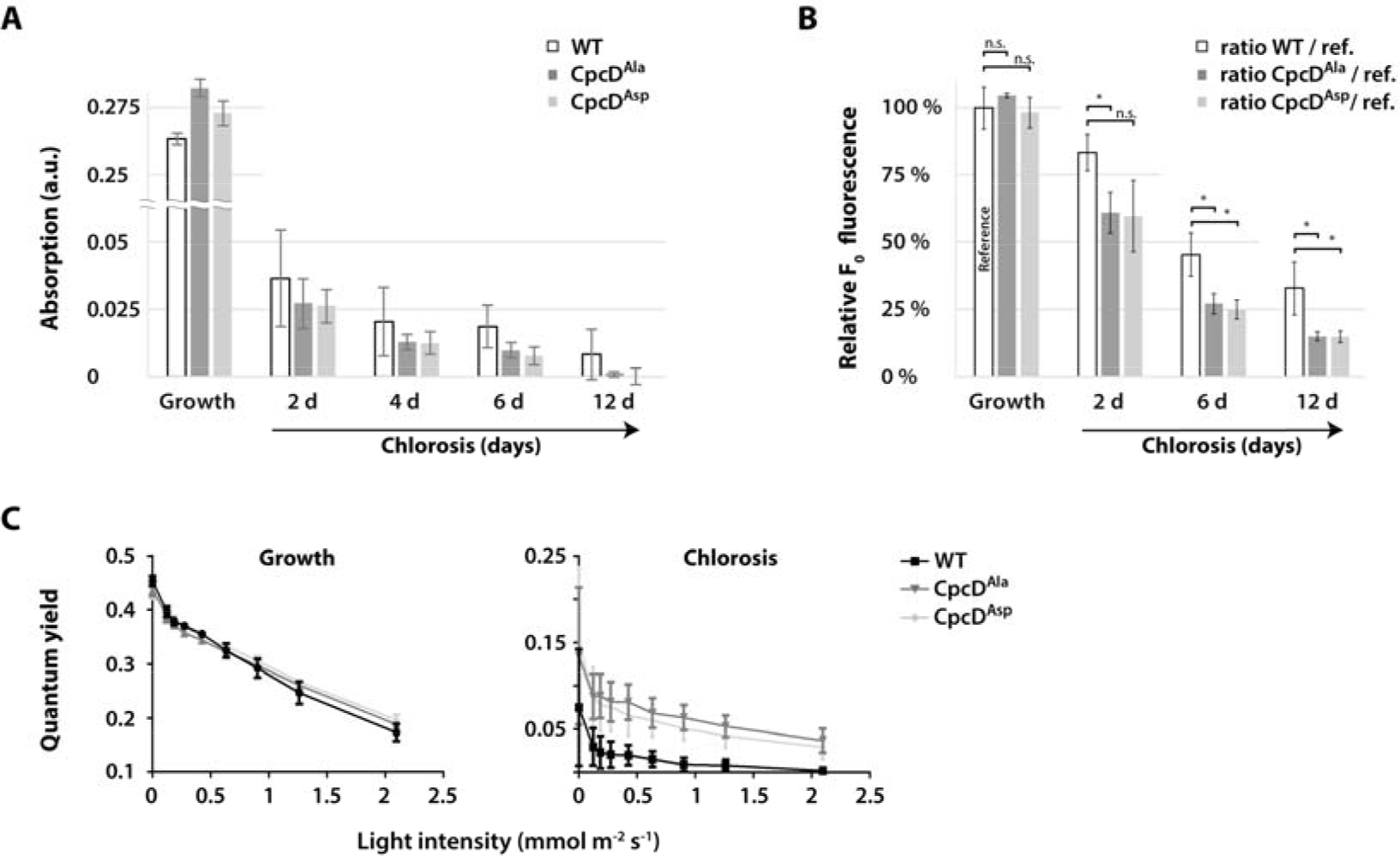
Phycobilisome turnover in CpcD mutants during chlorosis. **(A)** Comparison of phycobilisome absorption (625 nm) in CpcD^Ala^ or CpcD^Asp^ mutants and the wild-type during vegetative growth and during chlorosis. Shown are average values from three independent replicates at light intensities below 25 μM photons m^−2^ s^−1^ during the progression of chlorosis. Error bars indicate standard error of mean. **(B)** Determination of minimum fluorescence (F_0_) ratios of CpcD mutants and the wild-type during chlorosis relative to that during vegetative growth (reference). F_0_ fluorescence yields were recorded during chlorosis by PAM fluorometry of three independent replicates (the same cultures as in panel A). Ratios of average values to the reference (wild-type F_0_ during vegetative growth; defined as 100%) are indicated. Error bars indicate standard error of mean. Significant differences (p ≤ 0.05) between the wild-type and mutants are indicated by *; n.s. indicates non-significant differences. **(C)** Measurement of the PSII-phycobilisome quantum yield by PAM fluorometry at increasing light intensities during exponential growth and after 12 d of chlorosis. Error bars indicate standard error of mean, n = 3.

We used PAM fluorometry to investigate PSII activity in wild-type and mutant cells in more detail. In cyanobacteria, the basal PAM fluorescence signal (F_0_) results from overlapping PSII Chl *a* and phycobilisome fluorescence (38). In nitrogen-sufficient cells, the F_0_ levels of the mutants and wild-type were indistinguishable, but upon nitrogen depletion, F0 of the mutants was significantly lower than that of the wild-type (Figure 6B), which is in agreement with the lower phycobiliprotein level of the mutants. PAM light curves were used to characterize the light-harvesting properties (Figure 6C). In this assay, PSII quantum yield is recorded at increasing actinic light intensities. The PSII quantum yield of the mutants and the wild-type in the vegetative state was the same at all light intensities, which demonstrated that the CpcD mutations did not affect light harvesting in growing cells. However, in the chlorotic state, the quantum yield of the wild-type decreased much more strongly at higher actinic light than that of the mutants. The lower quantum yield is due to the closure of PSII reaction centers, caused by excess excitation. The maintenance of a high quantum yield in high light is characteristic of an inefficient light-harvesting system.

To further investigate the consequences of the CpcD mutations on the composition of the entire photosynthetic apparatus, we quantitatively compared the proteomes of the mutants and wild-type growing vegetatively and in chlorosis (Figure 7). The only marked difference between the mutant and wild-type proteomes of vegetative cells was the abundance of CpcD. The amount of CpcD in both mutants was about 22% of that of the wild-type; the differences in CpcD amount did not affect the abundance of other phycobiliproteins or the light-harvesting properties (see above). By contrast, all phycobilisome components were strongly reduced in the mutants in chlorosis. This result clearly confirmed our initial hypothesis that serine-phosphorylation of the terminal linker CpcD contributes to stabilization of the phycobilisome during chlorosis.

**Figure 7:**
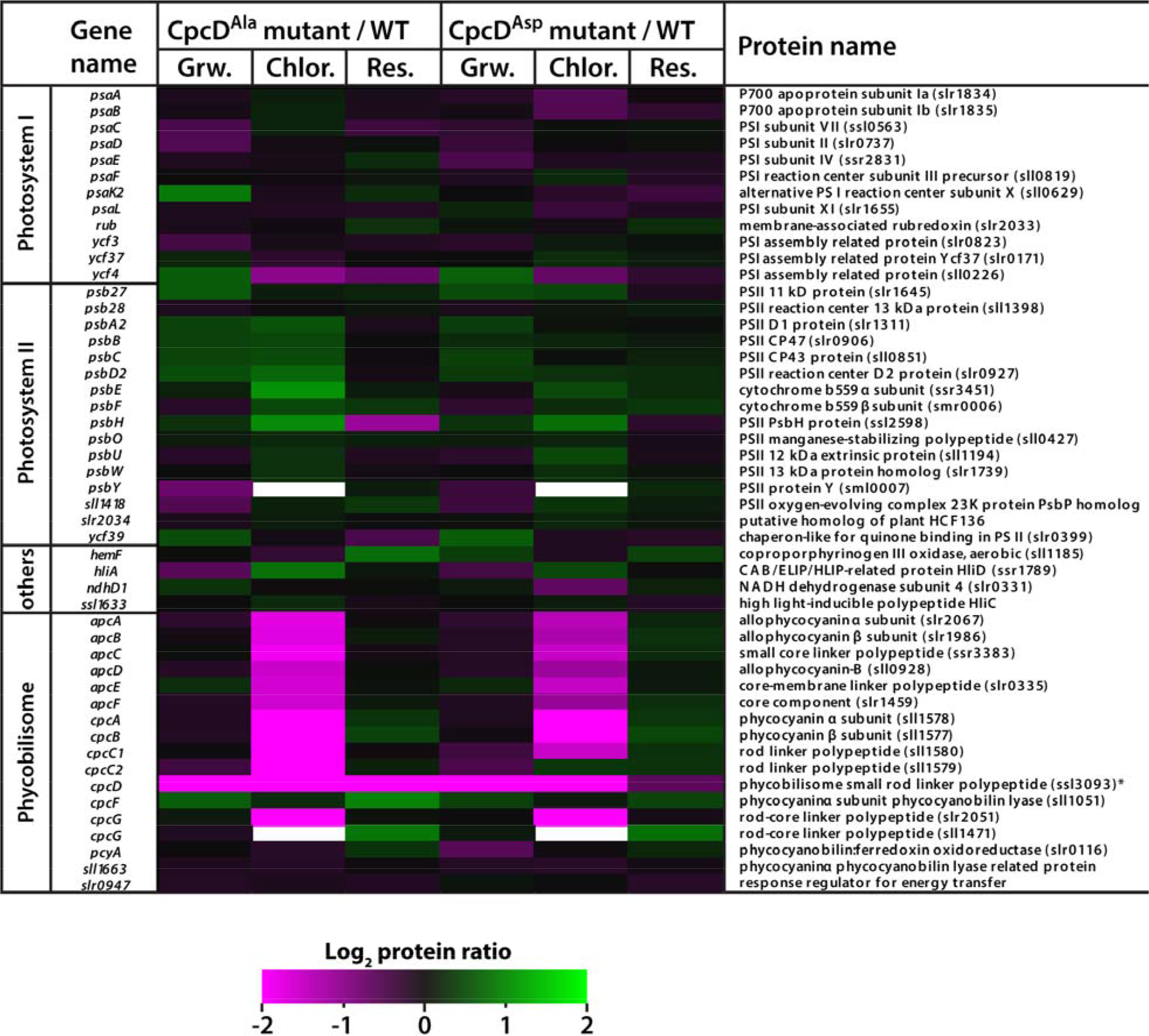
Quantitative comparison of the photosynthetic apparatus in the proteomes of CpcD^Ala^ and CpcD^Asp^ mutants and the wild-type. Proteins from cultures grown vegetatively (Grw.), in chlorosis for 6 d (Chlor.), or recovered from chlorosis (Res.; 48 h resuscitation) were extracted, digested with trypsin, dimethylated, and analyzed by mass spectroscopy. The relative abundances of the peptides of the mutants and wild-type were compared. Quantified peptide ratios (log_2_) of photosynthetic proteins were visualized in a heatmap (left panel: protein ratio CpcD^Ala^/WT; right panel: protein ratio CpcD^Asp^/WT). Proteins with lower abundance in CpcD mutants (protein ratio ≤ 0) are indicated by magenta, and proteins with higher abundance in CpcD mutants (protein ratio ≥ 0) are indicated by green.

We then used 77K fluorescence spectroscopy to functionally characterize the status of the light-harvesting system in the CpcD mutants. Again, during vegetative growth, the mutants did not differ from the wild-type (not shown). Chlorotic mutant cells only slightly differed from chlorotic wild-type cells following Chl *a* excitation (440 nm), which is in agreement with the slightly elevated Chl *a* concentrations in the mutants. By contrast, phycobiliprotein excitation (580 nm) of chlorotic cells yielded clearly different spectra in the chlorotic state (Figure 8). With wild-type cells, specific peaks from phycocyanin and allophycocyanin fluorescence (645 and 662 nm, respectively) were detected, but with mutant cells, no phycobilisome fluorescence was detected. These results unequivocally indicate that both CpcD mutants (CpcD^Ala^ and CpcD^Asp^) are unable to maintain functional phycobilisome complexes in the chlorotic state, highlighting the importance of CpcD for controlling the degradation of the phycobilisome under stress conditions.

**Figure 8:**
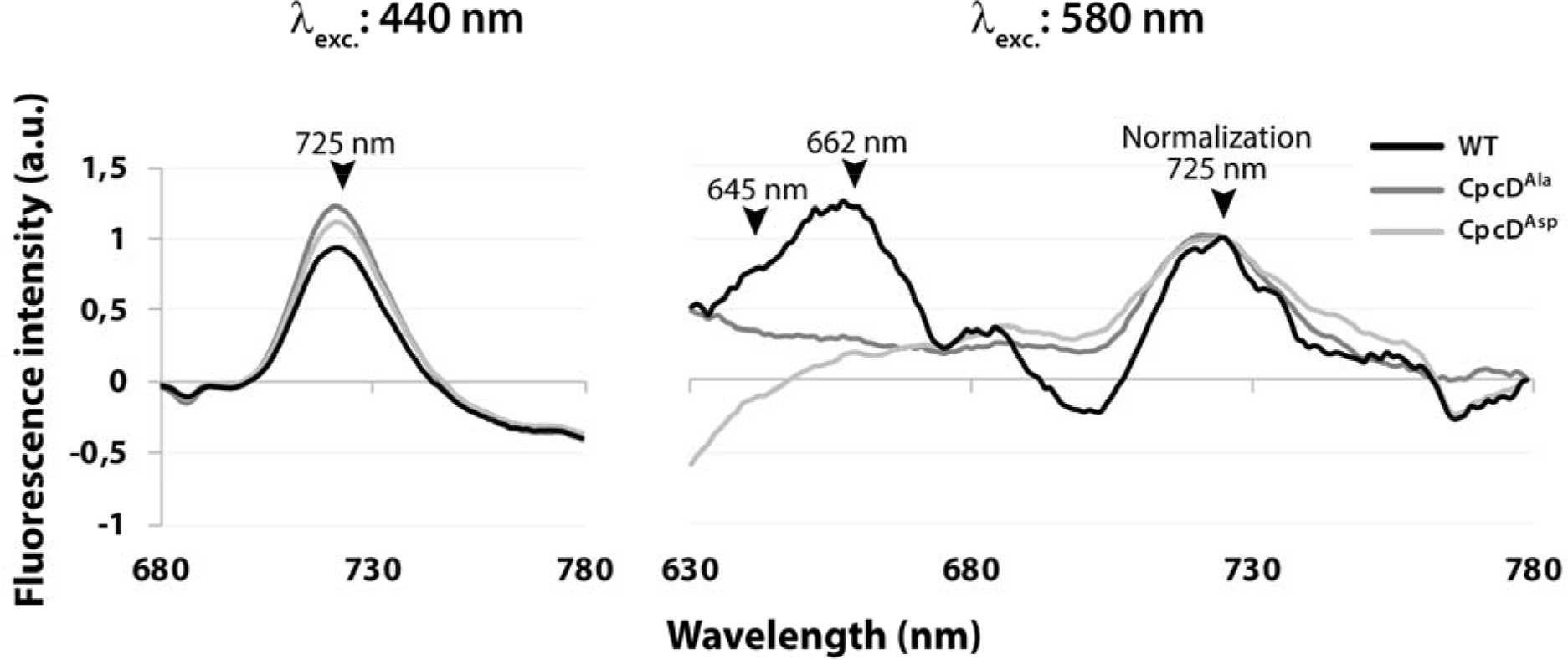
77K fluorescence spectroscopy of CpcD^Ala^ and CpcD^Asp^ mutants and the wild-type. Fluorescence emission of 21 d chlorotic cells was recorded **(A)** after Chl *a* excitation (440 nm) or **(B)** after phycobilisome excitation (580 nm). Maximum fluorescence signals of photosynthetic pigments are indicated in the spectra (PSI-Chl a: 725 nm; phycocyanin: 645 nm; allophycocyanin: 662 nm) and emission spectra from 580 nm excitation are overlaid (normalization) based on the 725 nm peak.

## Discussion

### Dimethylation Labeling Enables Deep Analysis of (Phospho)Proteome Dynamics in Cyanobacteria

Initial proteomic studies of (cyano)bacteria were limited to qualitative analyses. With the development of high-accuracy mass spectrometry and the steadily progressing sequencing power of nanoLC-MS/MS systems, deeper analysis of proteomes and their post-translational modifications became available (39). Today, gel-free sample preparations together with stable isotope labeling strategies set the standard for studying dynamic responses of the proteome and its post-translational modifications on global levels. Following this, the application of isobaric tags for relative and absolute quantitation (iTRAQ) represents the most common labeling strategy in cyanobacterial proteome studies (40), and references in (41). By applying dimethylation labeling as an alternative quantitation strategy (26) in our previous study, we enhanced the global proteome quantitation substantially, and, for the first time, obtained comprehensive O-phosphorylation dynamics (20). The high sensitivity of our improved workflow is of particular importance for long-term starved cells because the proteome is strongly degraded when cells enter a dormant state. In the current study, we quantified the dynamics of 2,271 proteins and 126 p-events with high confidence, which provides novel insights into the intriguing regulation mechanisms for cell survival and resuscitation. The reliability of our dataset is demonstrated by the striking congruence in the abundance of components of the rod and core parts of the phycobilisome, which perfectly matches the quantification by various biophysical methods, such as absorption spectroscopy and 77K fluorescence of the pigment proteins.

### The Status of Ribosomes in Chlorosis

Protein synthesis in long-term chlorotic cells is diminished to a barely detectable level (9), and electron micrographs reveal an almost complete absence of ribosomes in long-term chlorotic cells (11). At the level of the proteome of chlorotic cells, the least abundant ribosomal proteins set the upper limit for the abundance of assembled ribosomes. The presence of some ribosomal proteins in higher abundances is unexpected - if ribosomal proteins were completely degraded following disassembly of functional ribosomes, one would expect a similar abundance of all 56 ribosomal proteins. The highly divergent abundances of the different proteins indicates that this is not the case, and that some are not degraded to completion upon disassembly of functional ribosomes. Interestingly, the product of *ssl0438* (annotated as “similar to 50S ribosomal protein L12”) even accumulates in the chlorotic state (T0:T55 = 2.16), whereas authentic L12 is one of the ribosomal proteins with the lowest abundance. L12 composes the stalk of the large subunit, which is responsible for recruitment of translation factors. L12 is the only ribosomal protein present in multiple copies per ribosome (6 copies in *Synechocystis* sp.) (42). It is tempting to speculate that the relative increase of Ssl0438 indicates that the composition of the ribosomes in the chlorotic state is subtly modified as compared to those in vegetative cells. In support of this speculation, we found differential phosphorylation of several ribosomal proteins, including L12, in vegetative and chlorotic cells. Even more compelling, L9, which is the least abundant ribosomal protein in chlorosis, displays an 8.4-fold higher phosphorylation during chlorosis. Since L9 plays a key role in the fidelity of translation (43), its phosphorylation could represent a control point for overall ribosomal activity during chlorosis. In fact, a recent study has shown that L9 phosphorylation in *E. coli* helps survival during periods of starvation (43). Our results and earlier results together indicate that most ribosomes in chlorosis are disassembled, but not all ribosomal proteins are congruently degraded, and that the remaining functional ribosomes appear to be modified. During recovery, non-functional ribosomal proteins could be gradually replaced by newly synthesized ribosomal proteins. Indeed, the assembly of new functional ribosomes is one of the earliest processes in the awakening process, which seems necessary to support the subsequent restoration of cellular functions.

As a test case for the identification of potential unidentified components of protein complexes, we compared the expression profiles of hypothetical proteins in cluster 1 with those of ribosomal components, which are highly represented in this cluster. In this way, we identified the Ycf65-like protein (Slr0923), and the corresponding *slr0923* gene was clearly co-regulated with ribosomal genes at the transcriptome level (Supplementary Figure 9). Interestingly, Slr0923 contains a domain with sequence similarity to plastid-specific ribosomal proteins (PSRP domains) in plants; it has been proposed that this domain promotes protein-protein interactions in the ribosomal 30S subunit (44). Therefore, Slr0923 is presumably a functional component of the *Synechocystis* sp. 30S ribosome. In a broader sense, this bioinformatic analysis strategy might enable the assignment of potential functions to many other hypothetical proteins. Such an application would be especially useful for organisms with high percentages of unknown proteins, such as *Synechocystis* sp., which has 52% or 55% hypothetical proteins in common databases (cyanobase and uniprot, respectively).

### RuBisCO Biogenesis

The components of the CO_2_-fixing apparatus, namely the carbon-concentrating mechanism (Ccm) proteins and RuBisCO (RbcLS) subunits, are highly repressed during chlorosis at the transcriptome level, and carboxysome structures were not detected in electron micrographs (11). However, at the proteome level, the abundances of the corresponding proteins were higher than expected from these features. It has been proposed that during chlorosis, carboxysome shell proteins are substantially utilized as nitrogen resources (45). However, the protein dynamics of RbcL reflected the results of an earlier study, in which its levels remained at up to 50% after four days of nitrogen starvation (46). The detected RuBisCO proteins might reside in an inactive, disassembled state, potentially as a nitrogen storage utilized during resuscitation, as this observation has parallels to the dual role of RuBisCO in plant senescence (47,48). The early induction of RuBisCO genes, as detected in our previous transcriptome analysis (11), agrees with the early accumulation of the RuBisCO chaperon Slr0011 (threefold increase from T0 to T2) in the proteome. Slr0011 is essential for assembly of functional RuBisCO complexes (49), and its early accumulation indicates the beginning of the assembly of RuBisCO complexes. Interestingly, we found abundant phosphorylation on seven sites of the carboxysome structural protein CcmM during chlorosis and early time points of resuscitation (Supplementary Table S8). Thus, phosphorylation of CcmM might play a role in controlling carboxysome function, with decreasing phosphorylation at the end of the resuscitation process corresponding to the active state.

### Metabolic Enzymes Involved in Chlorosis

Proteins that increase in abundance during chlorosis could have important functions for survival. We identified several metabolic enzymes among these proteins that are likely directly or indirectly associated with glycolysis. For example, the abundance of the hypothetical protein Slr0337 was 34-fold higher during chlorosis—the protein with the strongest increase. Slr0337 carries an intrinsic glycoside hydrolase (family 57) domain, which in known proteins specifically cleaves 1,4-α-D-glycosidic bonds and some subsequently attaches 1,4-α-D-glycan fragments to another position within carbohydrates. The presence of this domain suggests that Slr0337 is involved in carbohydrate metabolism, specifically as a debranching enzyme in glycogen degradation (50). We identified another hypothetical protein in the same cluster with a glycoside hydrolase family 57 domain that increases in abundance during chlorosis. This protein, Slr1535, also contains a family 38 domain, which specifies it as an α-mannosidase. These findings agree with glycogen metabolism being a key to the entire chlorosis and resuscitation process (11). We addressed the specific roles of carbon catabolic enzymes for the resuscitation of chlorotic cells in a parallel study (manuscript submitted). Supported by the present proteome data, that study concludes that chlorotic cells already prepare the enzymatic machinery for glycogen degradation during the accumulation of glycogen in response to nitrogen starvation.

Another highly enriched protein (~tenfold) during chlorosis is a hypothetical type 2 NAD(P)H dehydrogenase (Slr0851). Unlike type 1 NAD(P)H dehydrogenases, which in cyanobacteria preferentially utilize NADPH for the reduction of plastoquinone in the respiratory electron transport chain, type 2 NAD(P)H dehydrogenases do not seem to contribute in electron translocation to plastoquinone. Instead, they might function as
cyanobacterial redox sensors and are potentially involved in redox-state regulation of the plastoquinone pool (51). Interestingly, deletion of the gene encoding a homologous type 2 NAD(P)H dehydrogenase leads to strongly compromised autotrophic growth, with impaired sugar catabolism and elevated glycogen accumulation; during heterotrophic growth, its deletion severely retards growth (52). Elevated levels of Slr0851 might be part of the machinery prepared during chlorosis for efficient sugar catabolism when resuscitation starts.

### Central Regulators

A key for the initiation of resuscitation is the assimilation of nitrogen via the glutamine synthetase/glutamine oxoglutarate aminotransferase (GS-GOGAT) reaction. Glutamine synthetase represents the main control point in this pathway. To ensure maximal incorporation capacity when nitrogen sources become available to chlorotic cells, glutamine synthetase has to be fully active. Therefore, the inhibitory proteins that regulate the glutamine synthetase enzyme, IF7 and IF17, must be tuned down. Accordingly, repression of the corresponding *gif* genes *(ssl1911* and *sll1515)* is further enhanced by antisense RNAs (11,53), and we observed the same trend at the protein level: IF7 levels remained at their lowest until cells reached the vegetative state and IF17 levels showed an increase at T24, which indicated the first signs of nitrogen replenishment. Conversely to the levels of the glutamine synthetase inhibitory proteins, the levels of the nitrogen-starvation-specific glutamine synthetase GlnN (Slr0288) (54,55) were threefold higher in chlorotic cells and lower after completion of recovery.

Overall, cluster analysis revealed a high number of hypothetical proteins involved in the recovery from chlorosis. These proteins were most prevalent in clusters 4–6. Particularly striking are the proteins in cluster 4, which responded immediately to nitrogen replenishment. Among these fast-responding proteins are three histidine kinases from two-component systems: Hik8 (Slr0750), Hik11 (Slr1414), and Hik33 (Sll0698). The gene *hik11* is essential, but its product has not yet been characterized (56). Hik8 interacts with the circadian rhythm protein KaiC, thereby regulating transcription of genes encoding enzymes involved in carbohydrate metabolism in response to the day-night cycle. Remarkably, Hik8 is required in *Synechocystis* sp. for the catabolism of accumulated glycogen during heterotrophic (dark) growth (57), and overexpression of *hik8* in S. *elongatus* results in the inability of glycogen accumulation during light exposure (58). The early accumulation of Hik8 detected in our dataset could therefore trigger glycogen utilization during resuscitation. The *hik33* gene encodes the redox-sensitive protein NblS, which is involved in the regulation of a variety of cellular stresses. Among other functions, NblS represses the *nblA* gene, which encodes the protease adapter protein NblA involved in phycobilisome degradation (59–61). Since *nblA* expression is strongly tuned down during resuscitation (11), the early accumulation of NblS might be related to this task, but could also have additional regulatory functions in redox regulation and osmoregulation (62).

### Phycobilisome Degradation During Chlorosis

Phycobiliproteins are degraded in an NblA-mediated process (63). It was proposed that degradation is initiated at the peripheral rod antennae and progresses towards the thylakoid-membrane-associated core components (8,63,64). Similarly, a recent study based on the analysis of single cells reported that after 24 h of nitrogen depletion, fluorescence from the rods (phycocyanin) diminishes prior to that of the core 34 (allophycocyanin) (65). According to this sequential degradation, the abundances of the core components remains higher than those of the antennae components. This hypothesis is supported by results of our earlier study, which addressed phycobilisome degradation at an early phase of chlorosis. There, we showed that phycobilisome rods were already at 45–65% lower levels after 24 h of nitrogen deprivation, whereas core proteins were still present at initial levels (20). In final chlorotic state, this trend is even more pronounced, with the rod subunits decreasing approx. to a five-times lower level than the subunits of the core (**Supplementary Figure 13**).

### Resumption of Photosynthesis

Phycobiliproteins started to increase in abundance before other photosynthetic components increased. Proteins associated with chlorophyll synthesis (cluster 2), especially enzymes for Mg-protoporphyrin and tetrapyrrole formation, and cytochrome *b* accumulated steadily over time. Results of our earlier transcriptome study indicated a sophisticated regulation of Chl *a* biosynthesis during resuscitation (11). A comparison of our earlier transcriptome data (11) and the proteome data presented here (Figure 9) indicates that one control point is the coproporphyrinogen III oxidase reaction. In the entire proteome, the only protein involved in Chl *a* biosynthesis was the oxygen-dependent coproporphyrinogen III oxidase HemF; the oxygen-independent enzymes HemN1 and HemN2 were absent. This indicates that heme biosynthesis for respiratory enzymes and phycobilin biosynthesis depends on oxygen. This ensures that these metabolites are only formed under oxic conditions; oxygen is essential in the first phase of resuscitation because it depends on respiration. A second crucial step, the synthesis of chlorophyllide, is catalyzed by light-dependent and light-independent protochlorophyllide reductases (Por and ChlB, and ChlL and ChlN, respectively). On the proteome level, we only detected the light-dependent Por, which implies that chlorophyll synthesis is only possible under light. This explains the light dependency of the transition from the first, respiratory phase of resuscitation to the synthesis of photosynthetic pigments (unpublished data).

**Figure 9:**
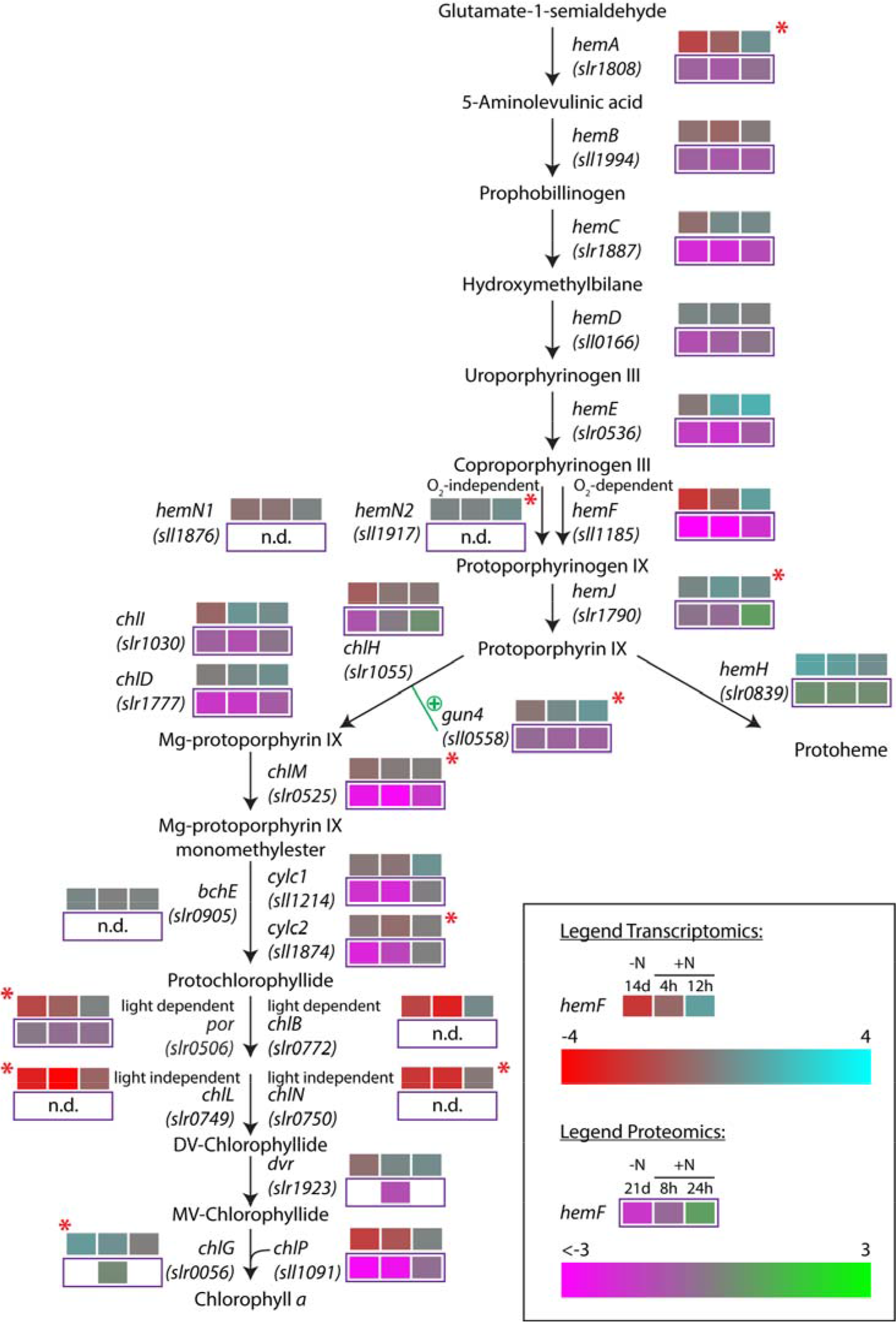
Regulation of the heme and Chl *a* biosynthesis pathway during resuscitation. Detected transcriptome and proteome dynamics of the involved enzymes are depicted for the respective time points of the resuscitation in a color code relative to vegetative and chlorotic conditions, respectively. Red stars indicate potential targets of antisense RNA regulation as detected by Klotz et al. (11).

Several components of the photosynthetic apparatus and associated F_o_F_1_-ATP synthase complex (PSII, PSI, and ATPase subunits) accumulated late in the resuscitation (cluster 3). In general, the corresponding transcripts were induced much earlier, especially those of ATPase subunits (11). Some of the early-expressed genes, such as those encoding ATPase proteins, might therefore initially replace non-functional copies of proteins that were not completely cleared in the chlorotic state. The net accumulation of ATPase subunits co-occurs with the synthesis of thylakoid membranes, where these protein complexes are embedded.

The abundance of the electron carrier protein cytochrome c_553_ (Sll1796), which transfers electrons from the cytochrome *b_6_f* complex to PSI, was low during chlorosis and only increased in the late phase of resuscitation. By contrast, the levels of the alternative electron carrier plastocyanin (Sll0199) (66) remained constant during chlorosis. Since both electron carriers participate in photosynthetic and respiratory electron transport chains (67,68), plastocyanin appears to be the dominant electron carrier in the chlorotic state.

Furthermore, the levels of two of three subunits of the electron-accepting cytochrome *c* oxidase (COX) complex (Slr1137 and Slr1136) were higher during chlorosis and early resuscitation. These enzymes could play an important role in maintaining redox homeostasis in the chlorotic state, together with the type 2 NAD(P)H dehydrogenase, whose levels were also higher. The respiratory activity of the COX reaction is also essential for the initiation of resuscitation. Therefore, maintenance of this complex appears to be a reasonable strategy to ensure rapid escape from chlorosis.

### O-Phosphorylation of Photosynthetic Components

The photosynthetic apparatus has always been a focus of cyanobacterial research. Not only the proteome, but also post-translational modifications in strains of *S. elongatus* and *Synechocystis* sp. have been investigated, and the occurrence of protein phosphorylation was already described in early studies of thylakoid membrane proteins (12,69). Likewise, the phosphorylation of a phycobiliprotein, later on identified as CpcB (70,71), in response to light quality and associated with state transition was reported early on (13,72). With shotgun-MS analysis, qualitative studies of cyanobacterial phosphoproteomes extended the knowledge about O-phosphorylation on the photosynthetic apparatus (18,19). Furthermore, the first targeted-MS analysis of photosystem components addressed the dynamics of selected phosphorylation events (37). However, the molecular function of O-phosphorylation remains widely unresolved (41).

The present study provides novel insights into the function of O-phosphorylation of photosynthetic components. While phosphorylation of PSI and PSII proteins is modestly dynamic during resuscitation, multiple phosphorylation events occur on two different peptides of the Vipp1 thylakoid membrane protein (cluster D). One of these peptides carried two phosphorylation sites, and the other carried six sites. However, only one of the phosphorylation sites of each of these two peptides was occupied at a time. This implies that the phosphorylation of any particular residue might prohibit further phosphorylation on neighboring sites. The phosphorylation level of most of these sites remained high until T24. The subsequent decrease occurs concomitantly with the accumulation of the Vipp1 protein and the re-emergence of a stacked thylakoid membrane, in whose biogenesis Vipp1 seems to be essential (73). In this process, Vipp1 forms large ring-like complexes associated with the thylakoid membrane (74). Phosphorylation might prevent Vipp1 complex formation during the first resuscitation phase. The appearance of unphosphorylated Vipp1 might initiate thylakoid membrane synthesis in the second resuscitation phase.

We detected extreme phosphorylation dynamics of several phycobilisome components. In particular, the phosphorylation sites of CpcA were close to 90% occupied in T0 and T2, and dropped to 6–24% in T55 (Table 2). Two phosphorylation events on CpcB showed an even more pronounced trend, with occupancies close to 100% in T0 and a rapid decrease to a final ~2–12% in T55. CpcB phosphorylation was previously associated with both phycobilisome state transition (13) and energy transfer in the phycobilisome (18). The residual phycobilisomes in chlorotic cells, which contain hyperphosphorylated CpcAB and CpcD proteins, seem to be resistant to NblA-mediated degradation. This assumption is in accordance with the finding that CpcB interacts with NblA (75), which triggers the Clp protease-mediated phycobilisome degradation process (76). However, CpcAB phosphorylation appears not to be sufficient to protect the phycobilisome from degradation, as evidenced by the almost complete degradation in the CpcD mutants. It is, therefore, possible that CpcAB phosphorylation is related to a starvation-specific state transition, in agreement with a previous study showing that state 1 to state 2 transitions are affected by starvation (77). The allophycocyanin core proteins, which represent the second target of NblA (63), are likewise phosphorylated, and ApcB had dynamics similar to that of CpcAB subunits.

### CpcD Phosphorylation Regulates Phycobilisome Turnover During Chlorosis

The *Synechocystis* sp. phycobilisome complex is generally composed of the allophycocyanin core and the protruding rods (composed of several hexameric CpcAB discs). These chromophoric components are interconnected by specific internal linker proteins and are attached to the thylakoid membrane (78). The terminal linker CpcD is located on the apical tip of the rods, and regulates the antennae length by preventing further elongation (79). A previous study (80) suggested that most of the *Synechocystis* sp. phycobilisome linkers, except CpcD, are phosphorylated during vegetative growth and dephosphorylated upon phycobilisome degradation, which occurs under high-light stress or nitrogen depletion. By contrast, our previous (20) and present studies never detected O-phosphorylation of these linkers. Piven et al. (80) used qualitative y^32^P-ATP autoradiograms that were extremely sensitive in detecting O-phosphorylation, which were possibly able to detect linker phosphorylation at levels below the sensitivity of mass spectrometry. Furthermore, the autoradiograms were not normalized to the decreasing amount of phycobiliproteins, and therefore, the disappearing y^32^P label could have been caused by general phycobiliprotein degradation. In any case, this linker phosphorylation is likely marginal and not relevant for phycobilisome stability.

In contrast to the internal phycobilisome linkers, the terminal CpcD linker turned out to be a prominent phosphoprotein, and its phosphorylation dynamics and occupancies correlated with those of CpcAB subunits. In fact, CpcD phosphorylation might be even more complex, as it is assumed that CpcD is a protein tyrosine phosphatase substrate (81). In any case, the similar phosphorylation pattern of CpcAB and CpcD led us to hypothesize that the terminal linker CpcD might also contribute to the stabilization of the phycobilisome complex against NblA-mediated degradation. Mutation of the CpcD phosphorylation sites to Ala resulted in almost complete phycobilisome degradation in the chlorotic state, whereas no difference was observed in the vegetative state. Surprisingly, a similar phenotype was obtained by mutating these seryl residues to Asp, a potential phosphomimetic mutation. It appears that for CpcD, Asp does not mimic phosphoserine, perhaps owing to its lower electrostatic potential or its different steric properties.

Using quantitative proteome analysis, we unambiguously demonstrated that the CpcD^Ala^ and CpcD^Asp^ mutations are specifically affected in the degradation of phycobilisomes. Other components of the photosynthetic apparatus were not impaired, except for slightly upregulated PSII core proteins, which might represent a compensation for the decreased light-harvesting antennae. These results indicate that phosphorylation of CpcD is essential for regulation of phycobilisome turnover and stability during chlorosis. We propose that a low percentage of phycobilisome complexes with phosphorylated CpcD linkers, formed in the vegetative state, initially escape the NblA-mediated degradation process upon starvation-induced NblA expression. In the chlorotic state, ongoing degradation of non-phosphorylated phycobilisomes leads to high phosphorylation occupancies of CpcD and other phycobiliproteins. The residual hyperphosphorylated light-harvesting complexes are functional and contribute to the maintenance of low levels of photosynthesis in dormant *Synechocystis* sp.

In summary, this work describes the global proteomic fundament of a profound cellular differentiation process that allows survival and escape from a dormant state. In general, the proteome dynamics reflect the sequential activation of cellular functions observed at the transcriptome level (11), suggesting that in contrast to seed germination, proteins are synthesized form newly synthesized mRNA. The function of abundant mRNAs in the chlorotic state awaits further investigation. The present data provides a rich resource for future projects, e.g. addressing the function of prominent hypothetical proteins and represents a roadmap to unveil the molecular processes underlying the reconstitution of a vegetative cyanobacterial cell.

## Acknowledgements

We acknowledge Dr. Karsten Krug for great support in cluster analyses. We thank Selina Schwarzbach and Lucía Gordillo Pérez from the Interfaculty Institute for Microbiology and Infection Medicine, Tübingen, and Kay Wilschinsky and Joseph Keller from the Department of Plant Biochemistry, Bochum, for help with experiments. We are grateful to Karen Brune for critically reading this manuscript. This work was founded by the German Research Council (DFG) via the research training group GRK 1708, the collaborative research center SFB 766 (A.K. & K.F.) and the research group FOR 2092 (S.R.).

## Data Availability

All (phospho)proteome raw data acquired by mass spectrometry were deposited at the ProteomeXchange Consortium (http://proteomecentral.proteomexchange.org) via the PRIDE partner repository (82) under the identifier PXD008027.

## Author Contributions

P.S., A.K., and K.F. designed the study and analyzed data. P.S. and A.K. prepared *Synechocystis* sp. samples and generated mutants. P.S. performed experiments together with B.M. (MS-based proteomics), S.R. (77 K spectroscopy), and K.F. (PAM). P.S. and K.F. prepared the manuscript with input from all authors.

